# Nonparametric Bayesian Contextual Control: Integrating Automatisation and Prior Knowledge for Stable Adaptive Behaviour

**DOI:** 10.64898/2026.02.26.708143

**Authors:** Sia Hranova, Stefan Kiebel, Michael N. Smolka, Sarah Schwöbel

## Abstract

Humans have a remarkable ability to act efficiently and accurately in familiar situations while remaining flexible in novel circumstances. Nonparametric contextual inference has been proposed as a computational principle that can model how agents achieve flexible yet stable behaviour in dynamic and possibly unknown environments. However, it remains an open question how humans learn, deploy and reuse stable contextual task representations so efficiently. To address this question, we propose the nonparametric Bayesian Contextual Control (NP-BCC) model, which integrates nonparametric contextual learning with two well-established cognitive mechanisms: repetition-based automatisation and schema-like prior knowledge. These two mechanisms are assumed to support behavioural stability and facilitate novel task acquisition. Simulations in dynamic multi-armed bandit tasks of increasing difficulty illustrate how the NP-BCC can acquire and reuse contextual task representations, with the proposed mechanisms operating in the intended, functionally meaningful manner. Specifically, we show via simulations that automatisation not only enhances task performance but also stabilizes contextual inference and structure learning, while structured prior knowledge accelerates the acquisition of novel contexts. We discuss the implications of our findings for computational accounts of adaptive behaviour and contextual learning, and outline directions for future empirical work, including investigations of context-dependent behavioural dysregulation relevant to conditions such as substance use disorders.

**Author summary:** People are very good at repeating well-learned actions in familiar situations, but they can also quickly adjust their behaviour when circumstances change. How the brain balances stability and flexibility is still not fully understood. There is growing evidence that the brain organizes experience into different “contexts”, which are mental representations of encountered situations. Computational models based on this idea can in principle reproduce flexible behaviour, but they often become unstable in complex environments. To improve stability, we borrow two simple strategies from everyday human behaviour. First, people tend to repeat actions that have worked well before. Second, when facing something new, they often reuse strategies from similar past situations. Using simulations, we show that combining these strategies with context-based learning produces more reliable behaviour in the model. Prior experience helps the model understand new situations more quickly, while repeated actions help stabilise behaviour once a situation becomes familiar. Taken together, our findings show how such mechanisms can give rise to both flexible and stable behaviour in the model.

## Introduction

Humans are remarkably well equipped to act with high accuracy both in known situations as well as when encountering a novel task for the first time, even when demands switch at a moment’s notice. It is an open question in psychology and neuroscience how humans achieve such a striking balance between maintaining a stable commitment to the current task and the ability to flexibly switch to a new one when appropriate. Striking this balance between stability and flexibility is essential for human cognition [1]. This balance has been at the core of a wide range of fields ranging, for instance, from learning [2], decision-making and cognitive control [3, 4] to clinical disorders [5]. For example, disruptions of stability–flexibility trade-offs have been implicated in substance use disorders, where substance-related habitual behaviour can become maladaptively persistent [6–8]. Computational accounts that explicitly model this trade-off may therefore offer a principled framework for examining such disorders.

One promising computational principle that describes flexible behavioural adaptation is context inference [9–14]. In this view, the brain subdivides its experience into distinct contexts. This division allows the brain to learn task-specific models of the environment [15, 16] which indicate the rules, such as contingencies or reward probabilities, for the current task at hand. It can use these models to choose behaviour relevant to the current task, while ignoring irrelevant information. However, context is often implicit, i.e., not cued, and has to be inferred as a latent state [10, 17]. Crucially, this inference process would also enable the brain to infer its uncertainty about the current context, i.e. how certain the brain is that the current task is the appropriate one [9, 11, 14]. This uncertainty readily mediates the trade-off between stability and flexibility: when certainty about the current context is high, the brain focuses on this context and task behaviour is accurate. When certainty is low, the brain becomes more ready to switch to one of the memorized contexts or to a new one.

More broadly, the computational challenge is not only to explain how humans infer which task they are currently facing and how they manage uncertainty about their tasks, but also how they build, reuse, and stabilize internal task representations across repeated experience. How such representations are formed when task structure is initially unknown, and how they come to support efficient behaviour without sacrificing flexible adaptation remains an open problem. Rather than seeking a complete theory of these processes, our goal here is to develop a computational framework that allows these interacting processes to be studied within a single formal setting. In what follows we briefly review and motivate the key computational ingredients to be incorporated in such a model.

From this perspective, the context inference view is in line with Bayesian theories of brain function, such as predictive coding and active inference, which cast cognition as probabilistic inference under uncertainty [18–21]. In these frameworks, neuronal computations are typically described as hierarchical, with higher levels encoding latent states that provide context for interpreting lower-level observations. Hence, contexts can be understood as high-level latent states that constrain perception, learning and action.

However, the number and structure of such contexts in our environment are typically not constrained [11, 13]. To capture this, computational models have turned to nonparametric Bayesian approaches [9, 22, 23], which allow agents to infer not only which context is currently active but also to adaptively learn new contexts when existing ones no longer provide an adequate explanation of experience. This mechanism of latent state discovery is commonly referred to as structure learning. One influential example of this approach is the COntextual INference (COIN) model [9], a nonparametric Bayesian model of motor learning that demonstrated how contextual inference combined with structure learning can capture a wide range of motor learning phenomena. The COIN model can infer both the number of latent states and their transition dynamics. In the field of value-based decision making, [22] have proposed a nonparametric contextual reinforcement learning model which can infer the number of latent states in an environment, but unlike the COIN model cannot learn contextual transition dynamics.

While the contextual inference framework has yielded important insights into how humans flexibly switch between tasks, it is typically formulated as a normative framework that assume fully rational inference over latent contexts [9, 13]. From this perspective agents optimally infer contextual structure given the available data without incorporating biases, heuristics, or prior knowledge. In contrast, human cognition is known to rely on structured inductive biases and heuristic strategies that simplify inference, reduce computational demands, and facilitate rapid acquisition of novel tasks [24–29]. Embedding such mechanisms into computational models provides a principled way to explore how biologically plausible constraints shape learning and decision-making and can therefore yield faster learning and higher overall performance than would be expected under purely normative models of contextual inference.

One such prominent and evolutionarily ancient mechanism which is ubiquitous across animals and humans is behavioural automatisation and the learning of habits [29–31]. Humans automatise behaviour when a task is well known and the same behaviour has been repeated in this task. This allows for fast execution of previously successful behaviour, and provides stability in the case of a well-known environment. Indeed, a recent proposal by [14], the Bayesian contextual control model (BCC), showed that combining contextual inference with context-specific forward planning and automatisation can reproduce a wide range of experimental findings across instrumental task learning, task switching and cognitive control [14, 32]. However, the BCC model is parametric and assumes a fixed number of possible contexts, and does not therewith incorporate structure learning.

Another well-known cognitive mechanism that aids structure learning in a multifaceted way is schemas. Schemas organize prior knowledge to give structure to contexts [33]. Their general and modular nature allows for the transfer and rapid application of knowledge across disparate tasks and additionally aides integration of novel, related information [26, 33]. Hence, schemas allow new tasks to be more easily acquired when prior, structured knowledge provides a scaffold for what such tasks may look like. Therefore, schemas can help achieve one- or few-shot learning when encountering a novel task in a way that would be difficult for a naive purely rational agent.

In this work, we present a computational model that brings together contextual inference, automatisation, and schema-like priors to investigate their joint contribution to adaptive behaviour. To achieve this we extended the BCC model [14, 32], by i) making context learning nonparametric and ii) introducing schema-like template contexts which give structure to newly opened contexts. Using this novel nonparametric Bayesian contextual control (NP-BCC) model we demonstrate using simulations (i) flexible establishing of new contexts and switching between them, (ii) stabilization of context inference and enhancement of task performance by automatisation, and (iii) novel context acquisition acceleration by a schema mechanism. These demonstrations aim to provide proof-of-principle illustrations of the framework’s capabilities rather than an exhaustive account of contextual cognition. We discuss the implications of our results for computational theories of cognitive control, relating our findings to psychological theory and outlining future directions for empirical testing.

## Methods

In the following, we formally describe the proposed nonparametric Bayesian contextual control (NP-BCC) model. We first briefly sketch the main ideas behind Bayesian nonparametrics and recap the original Bayesian Contextual Control (BCC) framework. We then introduce the full nonparametric contextual structure learning model and present the inference scheme used for online belief updating and context discovery in detail. Finally, we describe the simulation environments used to evaluate model behaviour in dynamic decision-making tasks. Mathematical derivations and additional implementation details are provided in the Supplementary material.

### Bayesian nonparametrics: DPs, HDPs, and HDP-HMMs

The Dirichlet process (DP) can be thought of as a generalization of the Dirichlet Distribution or the Dirichlet Mixture Model to an infinite number of outcomes or mixture components respectively [34]. The DP can be derived as the limit of the following finite mixture model as we take the number of components to infinity [35, 36]. Let’s assume a mixture model with *K* components and component probabilities *β* distributed according to a symmetric Dirichlet distribution *β* ∼ Dir (*γ*_new_*/K*, …, *γ*_new_*/K*) parametrized by some positive scalar *γ*_new_*/K*. Each component *k* is associated with some parameter *ψ*_*k*_ and specified by some generic function *F* (*ψ*_*k*_). Component parameters *ψ*_*k*_ are distributed according to their conjugate prior distribution *H*. Under this generative model for each data point *x*_*i*_ we first pick a mixture component by sampling a component indicator variable *c*_*i*_ according to probabilities *β*. The data point is then generated from the corresponding mixture component 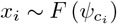. If we take the number of components in this model to infinity *K* → ∞, the resulting induced distribution *G* over component probabilities *β* and component parameters *ψ* is distributed according to a Dirichlet process *G* ∼ DP (*γ*_new_, *H*). The parameters of the DP are commonly referred to as the cluster opening tendency *γ*_new_ and the base measure *H*.

The DP can be used to perform conjugate posterior inference over both component parameters *ψ* and the number of components *K*. Importantly, in practice inference with the DP does not require representing an infinite number of components. Instead, posterior inference reduces to using a finite Dirichlet distribution over *K* known mixture components augmented with a special category that represents the probability of a previously unseen *K* + 1st component. This Dirichlet distribution is parametrized by the concentration parameters *γ* = {*γ*_1_, …, *γ*_*K*_, *γ*_new_} where the first *K* parameters can be interpreted as pseudo-counts which reflect how often each known component has been observed and are updated in the standard way. The *K* + 1st parameter is the cluster opening tendency of the DP and represents the probability of a novel component. Whenever a novel component is inferred, the dimensionality of the Dirichlet distribution is increased by one by adding a new parameter to the parameter set *γ* = {*γ*_1_, …, *γ*_*K*_, *γ*_*K*+1_, *γ*_new_}.

To model contextual transition dynamics nonparametrically, we use a Hierarchical Dirichlet Process Hidden Markov Model (HDP-HMM), which is a type of hierarchical DP. In this process a top-level DP defines a global set of possible contexts

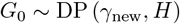

which induces a discrete distribution *G*_0_ over context probabilities *β* and parameters *ψ*. In the HDP-HMM the induced distribution over contexts *G*_0_ parametrizes a collection of lower-level or local DPs

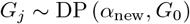

each of which defines a context-specific transition distribution. These local DPs correspond to the columns of a transition matrix *p* (*c*_*τ*_ |*c*_*τ*−1_) and induce the context-dependent transition probabilities *η*_*j*_ over the shared set of contexts. This construction of the prior over transition dynamics ensures that each column of the transition matrix is referring to the same set of possible contexts and enforces statistical regularization, with the mean of lower-level DPs being pushed towards the mean of the global DP [37]. Lastly, there is an additional self-transition bias parameter *κ* that pushes the expected *G*_*j*_ towards putting its whole probability mass onto the element *j*, thus making self-transitions highly likely [38].

### The BCC generative model

The NP-BCC is an extension of the Bayesian Contextual Control (BCC) model [14, 32], which can adaptively arbitrate between automatized and goal-driven control.

The BCC rests on the idea of *planning as inference* [39–41] which posits that an agent maintains a model of its knowledge about the world in the form of a probabilistic generative model, where actions are inferred alongside other hidden variables like future states and rewards. Because exact Bayesian inference in such models is analytically intractable, variational inference [42, 43] provides a practical approximation for estimating hidden variables. Along those lines, Active Inference (AINF) models describe value-based decision making and forward planning under the free energy principle [21, 44, 45] and offer a way to implement planning as inference while avoiding intractability. To model contextual processing using this approach [14] introduced a contextual layer within the active inference framework to account for context-dependent decision making, forward planning, and habit learning.

The resulting BCC model consists of two levels, a bottom level that describes forward planning and decision making on a per-context basis, and a top level which describes contexts and context inference on a predefined set of contexts. The bottom layer implements a context-dependent Markov decision process (MDP) and the top level describes contexts and their transitions as a Markov chain. Contextual transitions are assumed to evolve slower than the dynamics of the bottom MDP layer and select the rules active within a given behavioural episode. Once a context transition occurs and a new behavioural episode begins, the context remains fixed over the planning and decision-making horizon of the MDP. Taken together, the model uses a hierarchical partially observable Markov decision process (POMDP), which is defined by the tuple (𝒮, ℛ, 𝒜, 𝒞, 𝒯_*s*_, 𝒯_*r*_, 𝒯_*c*_), where

- 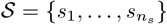 is a set of states
- 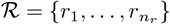 is a set of rewards
- 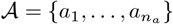 is a set of actions
- 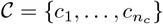 is a set of contexts
- 𝒯_*s*_ (*s*_*t*+1_|*s*_*t*_, *a*_*t*_) is a set of action-dependent state transition rules
- 𝒯_*r*_ (*r*_*t*_|*s*_*t*_, *c*_*τ*_)is a set of context-dependent reward generation rules
- 𝒯_*c*_ (*c*_*τ*+1_|*c*_*τ*_) is a set of context transition rules.

Note that here state transition and reward generation rules are treated separately as is typical in the active inference literature [21, 46], thus deviating from the notation used in the POMDP and reinforcement learning literature where rules are typically specified jointly [47, 48].

In line with the active inference literature, a behavioural policy *π* is defined as a deterministic sequence of actions *π* = {*a*_1_, …, *a*_*t*_, …, *a*_*T* −1_}. Then the POMDP can be formalized in a probabilistic generative model as

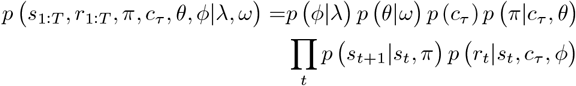

where

- *p* (*r*_*t*_|*s*_*t*_, *c*_*τ*_, *ϕ*) is a categorical distribution over rewards *r*_*t*_ parametrized by *ϕ* and conditioned on the current state *s*_*t*_ and context *c*_*τ*_ which the agent learns over time given by

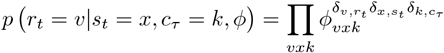

and *δ*_*i,j*_ is the Kronecker delta.
- *p* (*s*_*t*+1_|*s*_*t*_, *π*) is a categorical distribution over states *s*_*t*_ conditioned on the previous state *s*_*t*−1_ and an action specified by the policy *π*

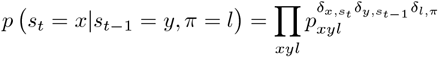
- *p* (*π*|*c*_*τ*_, *θ*) is a context-dependent prior over policies which the agent also learns over time and under which policies are distributed according to a categorical distribution parametrized by *θ* conditional on *c*_*τ*_

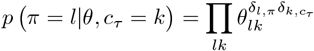
- *p*(*θ*|*ω*) is the conjugate prior distribution over the latent parameters *θ* of the prior over policies *p* (*π*|*c*_*τ*_, *θ*) which is a product of Dirichlet distributions

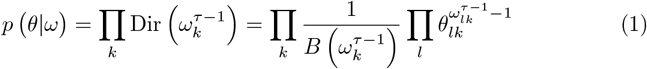

parametrized by the pseudo-counts *ω*^*τ*−1^ accumulated up to the previous episode *τ* − 1. Note this enables *ω* to be updated after each episode which enables the model to learn a prior over policies over time.
- *p* (*ϕ*) is the conjugate prior of over the latent parameters of the context- and state-dependent distribution over rewards *p* (*r*_*t*_|*s*_*t*_, *c*_*τ*_, *ϕ*) which is also a product of Dirichlet distributions

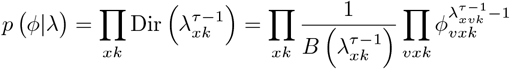

with pseudo-counts *λ*^*τ*−1^ accumulated up to the previous episode *τ* − 1. This allows the model to update *λ* and learn state-reward contingencies over time.
- *p*(*c*_*τ*_) is a prior predictive distribution over contexts with the transition probabilities averaged out.

Note that the BCC can be seen as a type of mixture model, where possible contexts correspond to mixture components and the component parameters for the *k*th component *ψ*_*k*_ = {*ϕ*_*k*_, *θ*_*k*_} become the MDP parameters {*ϕ*_*k*_, *θ*_*k*_} of the *k*th context. Importantly, mixture component memberships *c*_*i*_ represent the context that generated a given observation and are not independent and identically distributed. There is instead transition dynamics between mixture components *p* (*c*_*τ*_ |*c*_*τ*−1_) at each episode *τ*.

### The NP-BCC generative model

Because the BCC assumes a fixed set of contexts and a predefined transition structure, we extended it by introducing a nonparametric HDP prior over the context transition matrix *p* (*c*_*τ*_|*c*_*τ*−1_), following a similar approach to [9]. This prior enables the NP-BCC agent to learn the contextual transition structure of the environment in an unsupervised manner without having to predefine the number of contexts. To do this we developed an approximation to the HDP-HMM [37, 38] which enabled us to derive closed-form belief update equations for online variational inference. Here we present the resulting generative model of the NP-BCC.

In our approximation of the HDP-HMM we treated *G*_0_ and *G*_*j*_ as two independent constraints on the current context which are learned separately. As in the full HDP-HMM model, we introduce an additional context indicator variable *z*_*τ*_ associated with the global distribution over contexts *G*_0_. Together with the primary context indicator variable *c*_*τ*_, the variable *z*_*τ*_ is mapped to the corresponding context parameters *ψ*_*k*_ in their respective likelihood factors (see Eq (5) and Eq (6)). Unlike typical treatments of the HDP-HMM, we instantiate *z*_*τ*_ at each time point and link it to context parameters via a one-to-one mapping with *c*_*τ*_ rather than through nested indexing of parameters by indicator variables. Please refer to the Supplementary material for a detailed introduction to the HDP-HMM and our approximation.

The resulting prior does not allow for direct sharing of statistical strength and enforces a weaker regularization due to the breaking up of dependencies. Our approximation however retains essential HDP-HMM properties such as 1) being an infinite capacity prior over a transition matrix 2) *G*_*j*_ and *G*_0_ referring to the same set of contexts 3) inferring contexts based on a local and a global constraint; while allowing for closed form update equations for the HDP-HMM parameters *η*_*j*_, *β* and *ψ* (see Methods, Section Bayesian nonparametrics: DPs, HDPs, and HDP-HMMs).

The resulting NP-BCC generative model is graphically represented in Fig 1 and can be written down in full as

**Fig 1.**
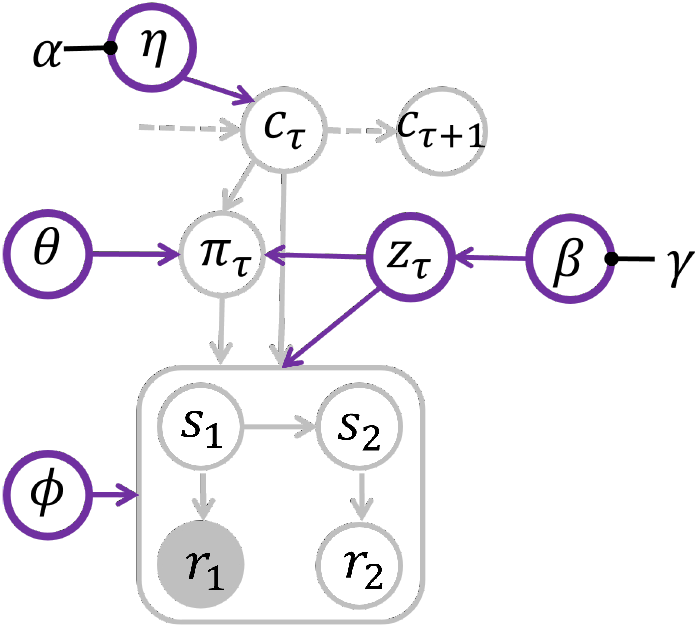
NP-BCC generative model as a Bayesian graphical model. This graphical model shows the NP-BCC generative model and conditional dependencies between variables. Empty circles represent latent variables and filled circles observed variables. Arrows represent statistical dependencies between connected variables and purple arrows represent novel or changed dependencies in the NP-BCC relative to the BCC. At the top level of the graph are the context indicator variables *c*_*τ*_ and *z*_*τ*_, while the lower level within the box contains the more rapidly evolving behavioural episode variables. At the beginning of episode *τ* the agent starts at state *s*_1_ and receives a reward *r*_1_. Knowledge about state-reward contingencies is represented by *ϕ*. In this graph the agent has already observed *r*_1_. It will then perform inference over future states and plan over possible policies, taking into account not only their goal-directed value but also the learned context-dependent prior over policies *θ*. It then executes an action which causes a state transition to *s*_2_ and the generation of another reward *r*_2_. At the end of an episode the agent updates its beliefs about reward generation rules *ϕ*, behavioural biases *θ*, context transition dynamics *η*, and context prevalence *β* by updating their associated hyperparameters (hyperparameters *λ* and *ω* not displayed for clarity). Note that at this point, the policy *π* has become an observed quantity. Then a context transition occurs from *c*_*τ*_ to *c*_*τ*+1_ and a new episode begins.

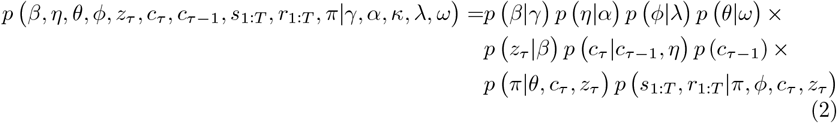

where

- *p* (*β*|*γ*) is the prior over context probabilities *β* introduced by the global *G*_0_ ∼ DP (*γ*_new_, *H*). This distribution is instantiated as a Dirichlet distribution over an expanding set of *K* possible contexts and a novel *K* + 1st context

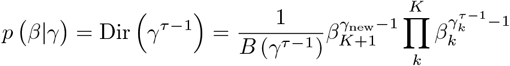

where *β* = {*β*_1_, …, *β*_*K*_, *β*_*K*+1_} specify the probability of the *K* known contexts and an additional *K* + 1st novel context. The concentration parameters *γ* = {*γ*_1_, …, *γ*_*K*_, *γ*_new_} encode posterior pseudo-counts 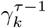 accumulated up to episode *τ* − 1 and *γ*_new_ which controls the prior tendency to instantiate new contexts and remains fixed.
- *p* (*z*_*τ*_|*β*) represents context assignment under the distribution *G*_0_ induced by the global DP. Under *G*_0_ contexts are categorically distributed according to *β*

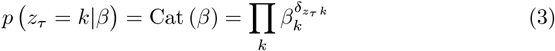
- *p*(*η*|*α, κ*) is the prior over context transition probabilities induced by the local DP (*α*_new_, *H*) and is similarly to above instantiated as

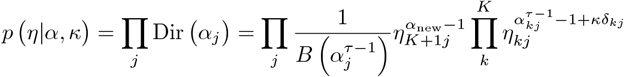

where *η*_*j*_ = {*η*_1*j*_, …, *η*_*Kj*_, *η*_*K*+1*j*_} and *α*_*j*_ = {*α*_1*j*_, …, *α*_*jj*_ + *κ*, …, *α*_*Kj*_, *α*_new_} with the the self-transition bias *κ* added to the diagonal of the pseudo-count matrix *α*. Please note that under our approximation of the HDP-HMM, global and local constraints on context identity are treated as independent, hence local DPs take the priors over parameters *H* = *p*(*θ*|*ω*)*p*(*ϕ*|*λ*) as a base measure rather than *G*_0_ as in the full HDP-HMM. For details please refer to the Supplementary material.
- *p*(*c*_*τ*_ |*c*_*τ*−1_, *η*) is the probability of going to a given context from a given context

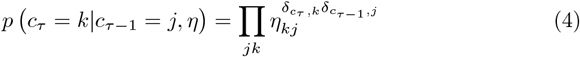
- *p*(*π*|*θ, c*_*τ*_, *z*_*τ*_) and *p* (*s*_1:*T*_, *r*_1:*T*_|*π, c*_*τ*_, *z*_*τ*_, *ϕ*) are the lower factors from the BCC (see Methods, Section The BCC generative model), where

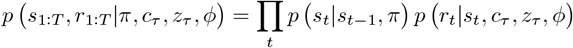

Note that in the NP-BCC all lower level context-dependent factors such as *p* (*π*|*θ, c*_*τ*_, *z*_*τ*_) and *p* (*s*_1:*T*_, *r*_1:*T*_ |*π, c*_*τ*_, *z*_*τ*_, *ϕ*) explicitly depend on both indicator variables *c*_*τ*_ and *z*_*τ*_. Similar to the full HDP-HMM, this is where the linkage between the global and local DPs happens, with the difference that in our approximation we have broken up the hierarchical dependency between the two and factors take the form of

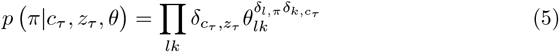

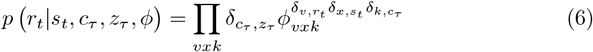

### Approximate Posterior

In order to infer the current context, future states, rewards and which policy to follow, the NP-BCC needs to calculate the posterior distribution over hidden variables

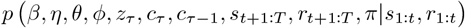

using variational Bayesian inference. Hyperparameters are left out for notational clarity. Posterior inference can be understood as the model asking: “given which states were visited, which rewards were received and which future rewards are desired, what is the most likely currently active context and its transition dynamics, what rules map states to rewards in this contexts, and what is the best course of action to reach desired states”. Bayesian inversion in the NP-BCC agent is carried out via variational inference. We factorize the approximated posterior *q* as follows:

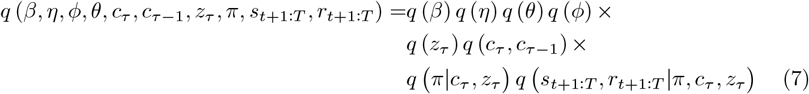

We used belief propagation message passing [49], see also [50], to infer the posterior over factors where variables critically depend on each other within or between episodes, like when inferring state transitions and state-reward rules *q* (*s*_*t*+1:*T*_, *r*_*t*+1:*T*_ |*π, c*_*τ*_, *z*_*τ*_) within an episode or context transitions *q* (*c*_*τ*_, *c*_*τ*−1_) between episodes. Other dependencies in the model were averaged out and a simpler mean-field approximation was used to infer the approximate posterior.

### Update Equations

The approximate posterior over the joint and marginal factors of the NP-BCC can be calculated at the minimum of the variational free energy [42, 49]

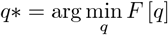

where the variational free energy *F* [*q*] is defined as

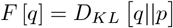

the Kullback-Leibler divergence between the generative model *p* (Eq (2)) and approximate posterior *q* (Eq (7)). These approximate posteriors represent the updated beliefs of the agent about hidden variables and parameters of the model. Here we present the relevant belief update equations.

#### Policy inference

At each time point *t* within an episode *τ* the NP-BCC perceives a state *s*_*t*_ and a reward *r*_*t*_. It uses this information to infer the probability of future states *s*_*t*+1:*T*_ and rewards *r*_*t*+1*t*:*T*_ on a per context and policy basis by calculating *q* (*s*_*t*+1:*T*_, *r*_*t*+1:*T*_|*π, c*_*τ*_, *z*_*τ*_) via belief propagation message passing (for details refer to [14, 32]). The agent uses this knowledge to plan into the future and infer a suitable policy based on its expected outcome. The posterior over policies

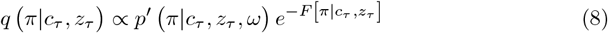

is proportional to the prior *p*^*′*^ (*π*|*c*_*τ*_, *z*_*τ*_, *ω*) times a likelihood in the form of the context- and policy-specific free energy

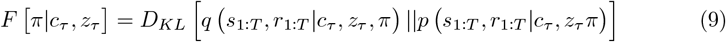

This free energy is lower, and the posterior is larger, when the predictions match the desired outcomes, i.e. a high likelihood of achieving rewards is predicted. This likelihood therefore encodes the goal-directed and value-based part of decision making in the NP-BCC. The prior *p*^*′*^ (*π*|*c*_*τ*_, *z*_*τ*_, *ω*) on the other hand encodes a priori biases to repeat policies that have been executed in the context before. This is enabled by the learning of the pseudo-counts *ω* in the conjugate prior (Eq (1)), which we averaged out in this equation according to the variational inference rules for notational brevity.

#### Context inference

The agent employs its updated policy beliefs to infer the posterior probability of the currently active context during the behavioural episode. As described in the previous section, there are two sets of indicator variables representing context due to the nonparametric prior, one stemming from the transition between contexts *p* (*c*_*τ*_ |*c*_*τ*−1_, *η*) (Eq (4)) and one from the global prior over contexts *p* (*z*_*τ*_ |*β*) (Eq (3)). The indicator variable *z*_*τ*_ is treated as auxiliary and used to calculate an approximate posterior over the currently active context *q* (*c*_*τ*_) that combines information from both the transition and global context probabilities.

The agent first calculates the joint marginal

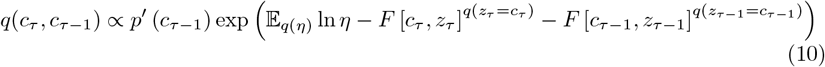

again via belief propagation message passing [42, 49]. Each term that makes up the posterior represents a message entering the factor *q* (*c*_*τ*_, *c*_*τ*−1_). The quantity *p*^*′*^ (*c*_*τ*−1_) denotes the message from the previous time point *τ* − 2 about the value of *q* (*c*_*τ*_, *c*_*τ*−1_) and is equal to 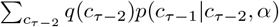. The quantity 𝔼_*q*(*η*)_ ln *η* is the message from the context transition factor with uncertainty over transition dynamics integrated out and 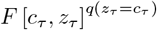 and 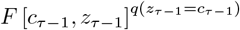 are the messages coming from the bottom layer of the graph which are weighted by the probability of a given context under the global prior *q* (*z*_*τ*_) as determined by our approximation. The approximate posterior *q* (*z*_*τ*_) is set to the predictive prior distribution *q* (*z*_*τ*_) = *p* (*z*_*τ*_ |*γ*^*τ*−1^)= ∫_*β*_ *p* (*z*_*τ*_ |*β*) *p* (*β*|*γ*^*τ*−1^) *dβ* once in the beginning of each episode.

The context-specific free energy evaluates to

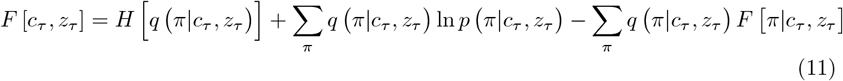

and *F* [*c*_*τ*−1_, *z*_*τ*−1_] is calculated analogously. Importantly, the context-specific free energy *F* [*c*_*τ*_, *z*_*τ*_] quantifies the agent’s approximate surprise associated with observed states, rewards and prospective actions for a given context. Contexts with lower free energy are assigned higher posterior probability. The three terms contribute to this quantity as follows.

The context-specific policy entropy *H* [*q* (*π*|*c*_*τ*_)] is lower the more certain the agent is about which policy will lead to favourable outcomes in a given context, which pushes the agent to infer contexts where it stays maximally uncertain about the best course of action, allowing for behavioural flexibility. The negative of the context-specific policy cross-entropy ∑_*π*_ *q* (*π*|*c*_*τ*_) ln *p* (*π*|*c*_*τ*_, *h*) is lower the more the posterior over policies fits to the prior over policies, i.e. it expresses how surprising the agent’s own behaviour is under the prior and favours contexts under which the executed policy is automatised. Lastly, the averaged out context- and policy-specific free energy ∑_*π*_ *q* (*π*|*c*_*τ*_) *F* (*π*|*c*_*τ*_) (Eq (9)) encodes how surprising the so far perceived reward-state pairs are given the agent’s knowledge about reward generation rules and favours contexts under which the observed outcomes are most likely. The final posterior over contexts is calculated by marginalizing the joint 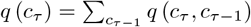 and is used for downstream policy selection and parameter update. Update equations for the model parameters *q* (*η*), *q* (*β*), *q* (*ϕ*) and *q* (*θ*) yield conjugate Dirichlet updates and can be found in the Supplementary material.

### Novel context opening

To perform online variational inference structure learning with streaming data and allow the capacity of the model to grow during inference, we employed the adapted mixture node proposed by [51] which emulates DP process structure learning.

In this mixture node the novel *K* + 1st context is treated as another mixture component associated with its own parameters *ψ*^novel^ = {*ϕ*^novel^, *θ*^novel^} which codify what a novel context looks like. In our implementation *ϕ*^novel^ and *θ*^novel^ were initialised to uniform distributions, encoding the assumption that a novel context is one where the agent knows nothing about reward generation rules and has no behavioural preferences. This allows for calculating the approximate surprise generated by the novel context *F* [*c*_*τ*_ = *K* + 1, *z*_*τ*_ = *K* + 1] and its posterior probability *q* (*c*_*τ*_ = *K* + 1) in the same way as for any other known context. If the novel context becomes the most likely explanation under the posterior *q* (*c*_*τ*_ = *K* + 1) ≥ 0.5, a new context is initialized and added to the set of possible models to perform inference over in the next episode by extending the capacity of *ψ, β* and *η* by one.

### Inference procedure

The resulting inference procedure and belief update scheduling is presented in Algorithm 1.

#### Algorithm 1

Online Variational Inference with DP Mixture Node

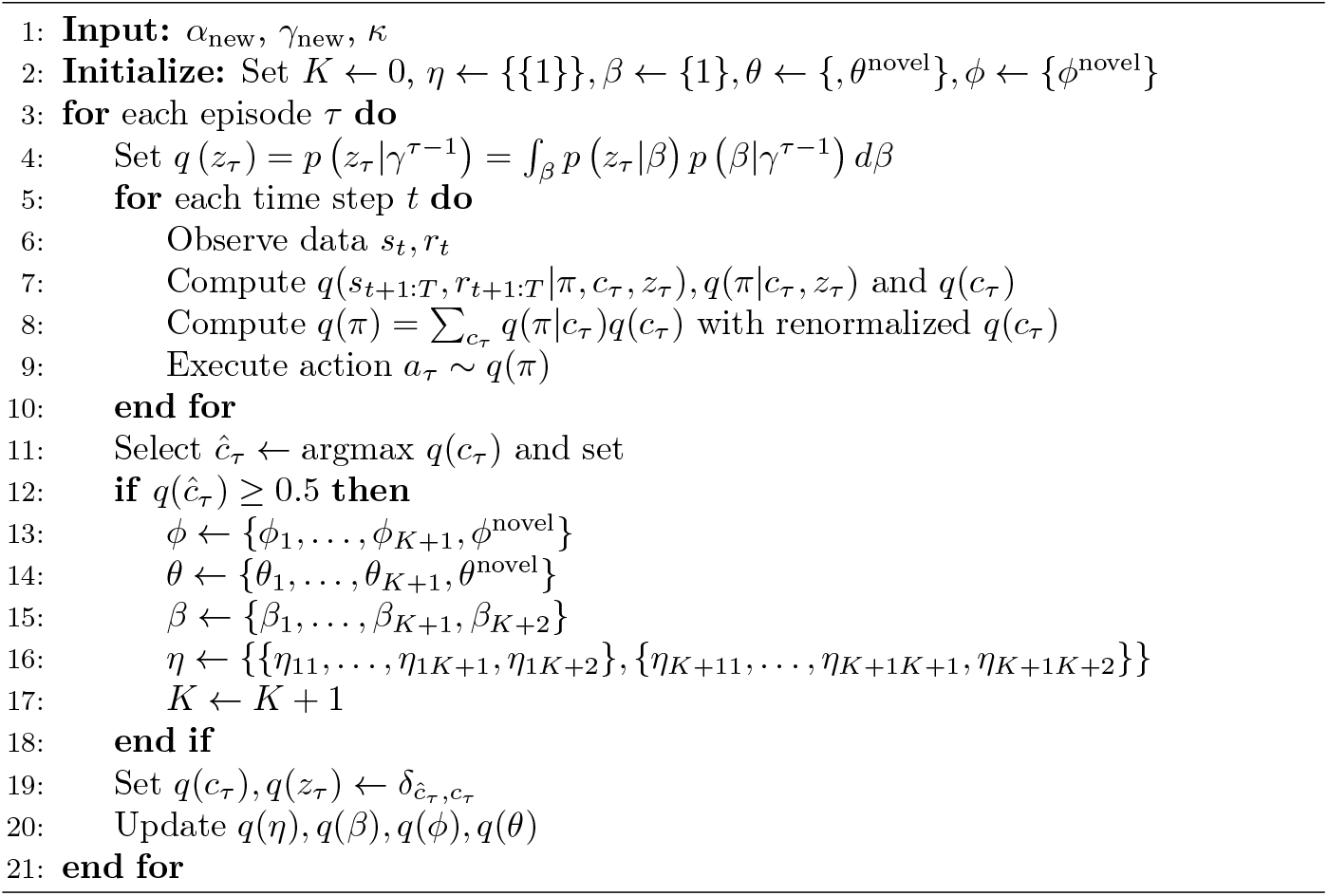

At each time point *t* within an episode *τ* the agent infers beliefs over future states *s*_*t*+1:*T*_ and rewards *r*_*t*+1:*T*_, context *c*_*τ*_ and policy *π* given received states, rewards and reward preferences. It then samples an action from a policy in proportion to the marginalised policy posterior 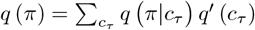, where *q*^*′*^ (*c*_*τ*_) is renormalized to exclude the probability of the novel context *q* (*c*_*τ*_ = *K* + 1) from *q* (*c*_*τ*_). At the end of an episode the agent makes a decision as to whether a new context has occurred based on its final context posterior estimate *q* (*c*_*τ*_) and expands model capacity by one if the novel context generated the least surprise, i.e. *q* (*c*_*τ*_ = *K* + 1) ≥ 0.5 (see Methods, Section Novel context opening). For numerical stability during online inference, newly opened contexts were initialized with a large global prior pseudo-count *γ*_new_ = 1000 to prevent early dynamics from being dominated by the auxiliary global indicator *q*(*z*) and to shift learning of persistence and switching onto the transition parameters *η* and their pseudo-counts *α*. Additionally, the transition matrix *η* was made square for inference purposes by adding *η*_*K*+1_ with a high self-transition bias.

Importantly, after a context opening decision is made, the inferred posterior over contexts is collapsed onto the maximum a posteriori context *ĉ*_*τ*_ = argmax*q*(*c*_*τ*_) to perform downstream parameter updates [51]. This is required for emulating the data point allocation procedure of the Chinese Restaurant Process formulation of the DP (see Supplementary material for explanation of the CRP).

Lastly, the agent updates its beliefs over model parameters such as context transition dynamics *η*, global prior parameters *β*, context-specific state-reward mapping rules *ϕ* and behavioural biases *θ* according to their respective update equations (see Supplementary material).

### Model setup and relevant hyperparameters

In the results we will show the influence of two key model components and their associated hyperparameters on context inference and behaviour: automatisation learning as represented by the automatisation tendency, which varies how quickly a prior over policies is learned, and context templates, which determine the parameters that are loaded into newly opened contexts.

#### Automatisation learning and automatisation tendency

As in [14], the NP-BCC is capable of learning automatisms in the form of an a priori tendency to repeat policies that have been chosen in the past. This is implemented in the context-specific prior over policies (see Eq. 8) where the pseudo-counts *ω* keep track of how often a policy has been executed in a given context. These pseudo-counts start from an initial value *ω*_init_, which determines how quickly the prior becomes sharp and starts dominating behaviour. To quantify such an automatization speed in a hyperparameter, we define the *automatisation tendency* as 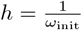. Here, an *h* close to zero means the prior stays almost constant and close to uniform, while an *h* close to one means that the prior is learned quickly and the bias can alter decision making within a few repetitions of the same policy.

#### Template contexts and reward hyperparameter initialization

In the NP-BCC we explored the effects of building in structured prior knowledge into newly opened contexts. In our simulations agents performed a multi-armed bandit multiple choice task (see Methods, Section Simulation environments). In such a task it is reasonable to assume there is a unique most rewarding choice option within a given context, as dictated by Occam’s razor. We built this assumption into the model via *template contexts* which represent an initial guess about what the state-reward contingencies of a newly opened context could look like. Once a template context is loaded in by initializing the contingencies of the novel context *ϕ*_*K*+1_ to those of the selected template, the agent proceeds to learn the true contingencies associated with this context.

We instantiated a set of template contexts 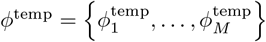, where 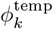 implies that the *k*th bandit is the most rewarding one. This is achieved by associating 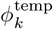 with initial pseudo-counts 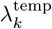 that assume that a reward has been received 11 times and no reward 1 time in state *k* and a reward was received 1 time and no reward 11 times in all other states. Pseudo-counts were chosen to encode a strong but non-deterministic prior preference and were selected for practical stability. They encode the assumption of a clearly optimal state and policy under the template contingencies 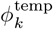 while allowing contextual contingencies to adapt within a few trials or, if the loaded assumption proves incorrect, the opening of a new context.

Templates were selected for loading based on which template generated the lowest prediction error with regards to the observation that caused context opening, whenever a new context was detected based on the uniform contingencies *ϕ*^novel^.

### Simulation environments

Agent performance was probed using a non-stationary multi-armed bandit (MAB) task. This task or a formal equivalent thereof is commonly used in the behavioural sciences to probe various aspects of decision making such as the arbitration between goal-driven and habitual behaviour or the the stability vs flexibility trade-off and others [22, 52–58]. In a MAB task agents are presented with *M* choices or bandits, each of which can lead to receiving or not receiving a reward with some probability. In a non-stationary MAB task, the true bandit reward contingencies can change. Agents must select bandits to gather as much reward as possible. Because bandits confer reward probabilistically, agents must infer the currently most rewarding bandit. We associate contexts with the different possible bandit reward contingencies (Fig 2).

**Fig 2.**
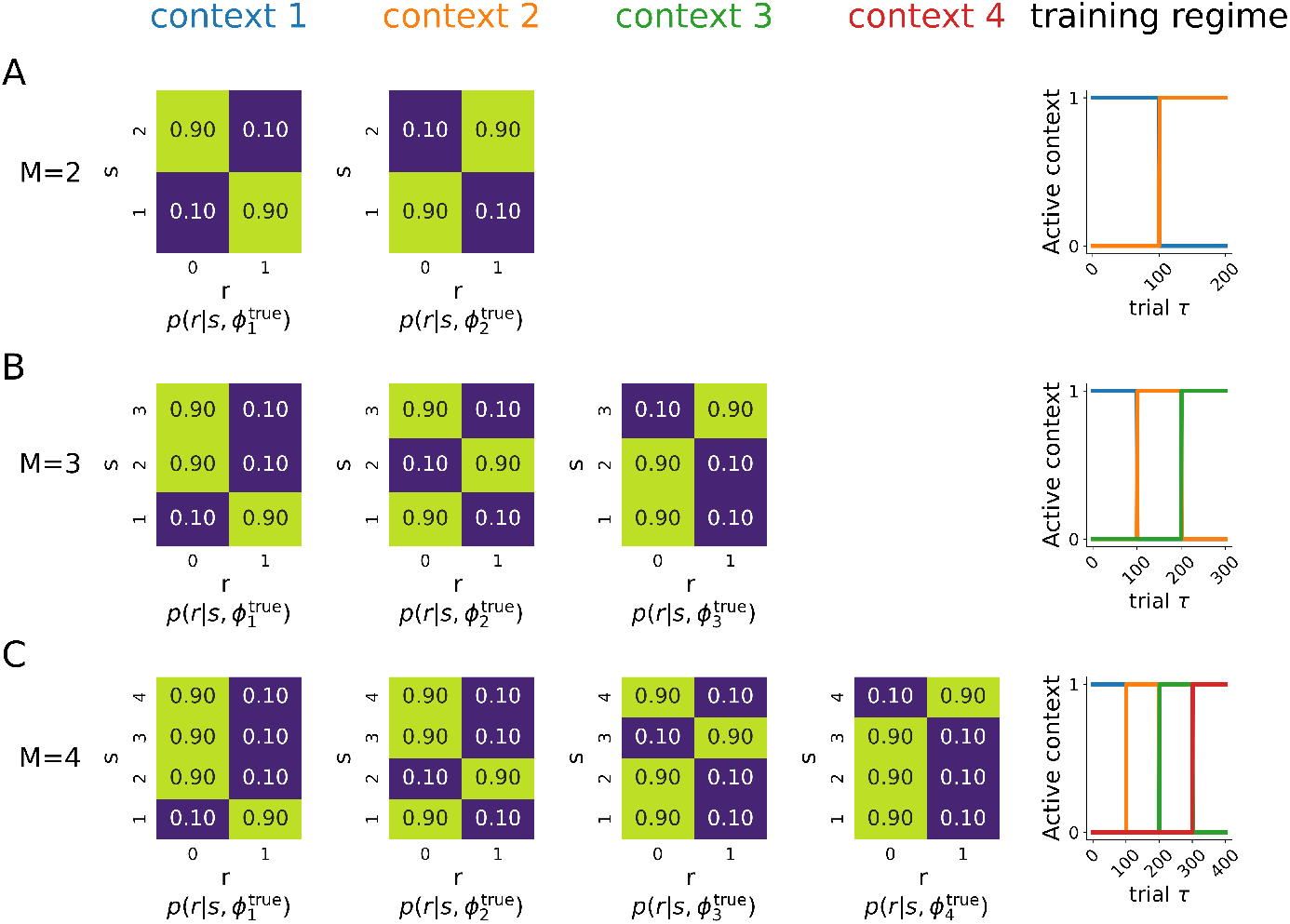
Simulation setups for multi-armed bandit (MAB) task. **A)** Panel shows simulation setup for a MAB task with *M* = 2 possible bandits. Matrices show the true state reward contingencies 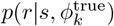 for the possible contexts in this task. State reward contingencies express the probability or receiving a reward *r*, given that the agent is in a state *s* of having selected a given bandit. These probabilities are parametrized by the respective parameters 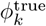. In the *M* = 2 task there are two possible contexts, as indicated by the two matrices: context 1 (associated with the blue colour) where bandit 1 is optimal with 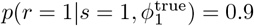 and context 2 (orange) where bandit 2 is optimal with 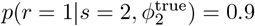. The training regime dictates which of the two possible contexts is active at a given trial. The active context switches every 100 trials. So agents first experience a 100 trials where (blue) context 1 is active and then another 100 where (orange) context 2 is active. **B)** Panel shows the simulation setup for a MAB task with *M* = 3 possible bandits. The setup is analogous for the *M* = 2 case with the difference that there is three bandits and correspondingly the environment has three possible contexts, a blue one where bandit 1 is optimal, orange one where bandit 2 is optimal and a green one, where bandit 3 is optimal. Training again consists of blocks of 100 trials, where one of the three possible contexts is active. **C)** The panel shows the setup for a MAB task with *M* = 4 bandits. Setup is analogous to above, with a fourth possible context added where bandit 4 is optimal (red).

The simulation environment is defined by (i) the number of possible bandits *M* (ii) the total number of possible contexts *K*^true^ which is the same as the number of possible bandit reward contingencies (iii) the true contingencies 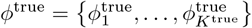 associated with each context which parametrize the probability of receiving a reward given that we are in the state of having selected one of the bandits 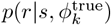 and (iv) a training regime which specifies which context is currently active, i.e., which contingency 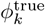 defines the probability of receiving a reward if we select a bandit at a given trial.

We simulated three different environments, with *M* = 2 (Fig 2A), *M* = 3 (Fig 2B) and *M* = 4 bandits (Fig 2C). We set up all simulations such that within a given context there is always an optimal bandit that would deliver reward with probability *p* = 0.9. All other bandits would deliver reward with only *p* = 0.1 in that context. Correspondingly, each context had a unique optimal action.

## Results

In this section we demonstrate that the proposed NP-BCC model can make adaptive decisions in uncertain and dynamic environments by using contextual uncertainty to regulate the trade-off between behavioural flexibility and stability. We begin with a simple binary choice task in which reward contingencies change abruptly and without cueing, using this toy example to illustrate the core inference and structure learning mechanisms of the model. This example shows how the NP-BCC can, in principle, acquire an expanding repertoire of context-specific memories through surprise-driven contextual structure learning.

We then turn to the limitations of such naive structure learning in more complex environments and show, through computational experiments, that performance degrades as task complexity increases. Finally, we demonstrate how task performance is stabilized and novel context acquisition is accelerated when two cognitive mechanisms that have so far been studied largely in isolation, repetition-based automatisation and the use of schema-like structured prior knowledge via template contexts are incorporated into the NP-BCC model.

### Didactic example: structure learning and contextual inference in the NP-BCC

In this section we present a didactic example of an instance of the NP-BCC agent performing a multi-armed bandit (MAB) task with *M* = 2 choice options. The example is meant to demonstrate how the model acquires and applies contextual representations of an environment via integrating contextualized value-based processing and structure learning. To keep things simple initially, we showcase a naive structure learning agent that does not automatize or deploy schema-like templates.

As described in Methods, Section “Simulation Environments”, in a *M* = 2 MAB task agents have to choose between two options called bandits, each of which can deliver a reward with some unknown probability (Fig 2A). In this environment, reward probabilities remain constant for 100 trials before switching abruptly without explicit cues. Each latent context is defined by a distinct set of reward probabilities over the two bandits. In the present simulation, there are two contexts: one in which bandit 1 yields reward with high probability (90%) and bandit 2 with low probability (10%), and a second context with the opposite contingency. Because the active context is not directly observable, the agent must infer both the current context and its contingencies from its action-reward history.

Structure learning is generally understood as the process of learning a hierarchical representation of an environment, where the complexity of the learned representation scales with the complexity of the observed data. Concretely, in this experiment structure learning means that an agent has to infer both how many contexts there are, and which reward probabilities *ϕ* they are associated with. The NP-BCC agent achieves this iteratively by deciding whether a qualitatively novel context has occurred or whether a known context can account for the recent observations. It does so by comparing the surprise elicited by observations and behaviour under the *K* contexts it already knows, to the surprise generated by a novel *K* + 1st context associated with uniform contingencies *ϕ*^novel^ (see Algorithm 1 and Methods, Section Novel context opening for details).

In the beginning of the simulation the agent begins with an empty set of contexts and *K* = 0. At the first trial the agent opens a context with probability 1. Initially the agent knows nothing about this context. Through trial and error it quickly learns the contingencies *ϕ*_1_ associated with context 1, i.e., that in the current context bandit 1 has a high chance of delivering reward (Fig 3A top). Correspondingly the agent infers that it is in context 1 (blue context in Fig 3B) as long as the reward probabilities stay constant and consistently executes the correct action of selecting bandit 1. This is because gathered observations are more likely under the contingencies *ϕ*_1_ than under the contingencies *ϕ*^novel^ associated with a novel context (Fig 3A top).

**Fig 3.**
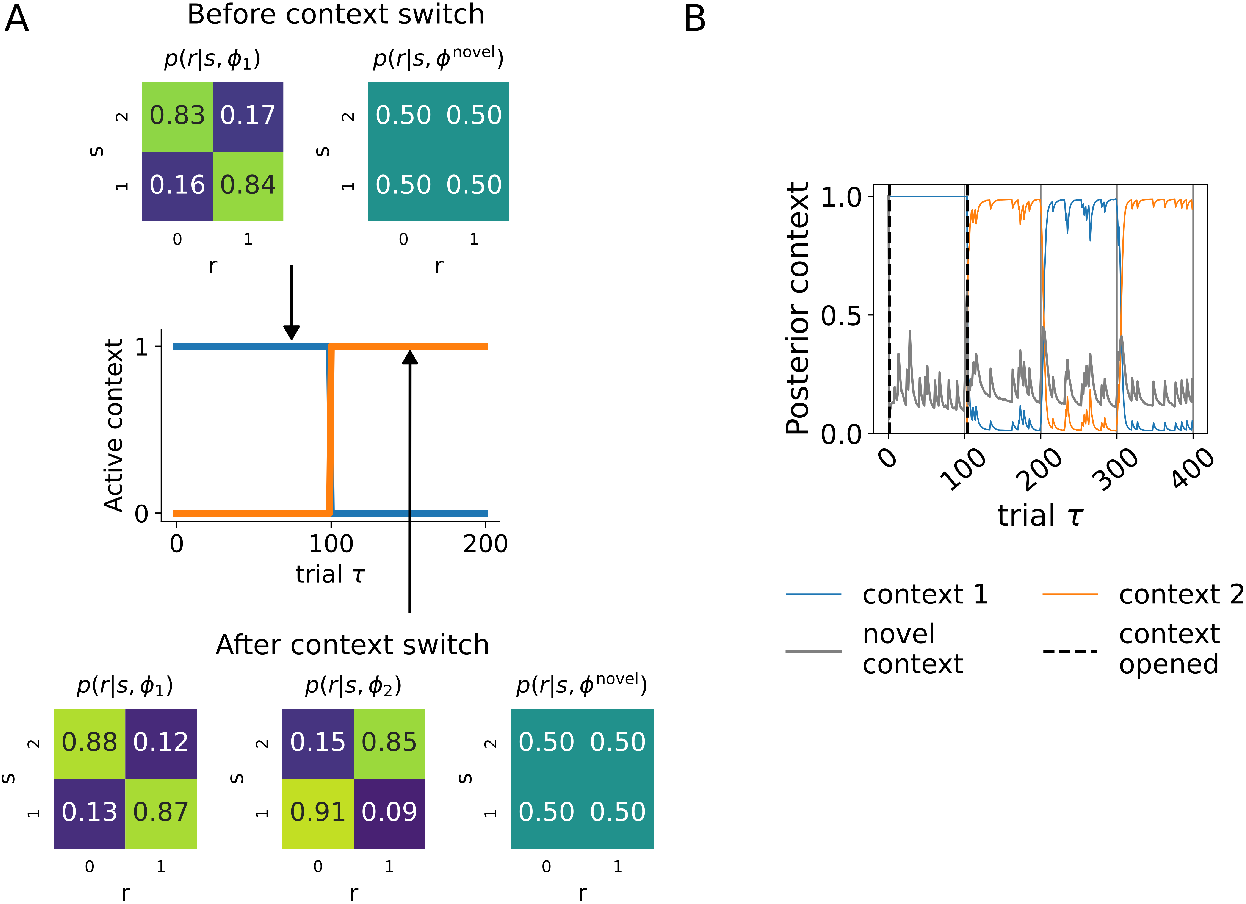
Contextual inference dynamics of an agent in the *M* = 2 MAB task. **A)** The panel depicts visually what happens when an agent opens a new context. The line plot in the middle of the panel shows an example training regime for the *M* = 2 MAB task, where in the first 100 trials the blue context associated with bandit 1 being optimal is active, and after a context switch at trial *τ* = 100 the orange context where bandit 2 is optimal becomes active. The matrices above and below the training regime plot represent the reward contingencies learned by the agent before and after a context switch has occurred. Each matrix encodes *p*(*r* | *s, ϕ*_*k*_) or the probability of receiving a reward *r* given that bandit *s* was selected under the reward contingencies *ϕ*_*k*_ associated with context *k*. Before the context switch (top row of matrices) the agent has experienced only the blue context and has correspondingly learned that bandit 1 has a high chance of delivering reward and bandit 2 a low chance of delivering reward as indicated by *p*(*r*|*s, ϕ*_1_). The agent additionally represents the possibility of a novel, yet unknown context defined by the parameters *ϕ*^novel^ which encode a uniform distribution of rewards given bandits. After the context switches at *τ* = 100, the orange context becomes active. The agent detects the presence of a novel context through the stacking surprise experienced under the contingencies *ϕ*_1_ associated with the blue context and extends the set of contingencies that it learns with *ϕ*_2_, while still representing the possibility of a yet another novel context via *ϕ*^novel^. With further training the agent correctly learns the contingencies associated with the orange context (bottom row of matrices). **B)** This panel depicts the inferred renormalised posterior over contexts *q*^*′*^(*c*_*τ*_) in colour (see Methods, Inference procedure) and the novel context probability *q*(*c*_*τ*_ = *K* + 1) in gray from a single instance of an agent performing a *M* = 2 MAB task. The agent experienced 100 trials of the blue context, followed by a 100 trials of the orange context. It then performed 100 trials of each context again. Vertical lines represent when a context change took place in the environment (solid vertical lines) and when the agent chose to open a new context (dashed vertical lines). The agent correctly inferred the presence of two contexts, shown in blue and orange, respectively. It additionally inferred each context with high certainty only in trials in which it was actually active. For clarity the posterior probability of possible active contexts is plotted after excluding the probability of a novel context occurring and renormalizing.

After the true context switches at *τ* = 100, bandit 2 becomes the optimal choice. At this point, the surprise generated by the previously learned contingencies *ϕ*_1_ of context 1 rises as the agent selects bandit 1 and fails to receive a reward. As a consequence, the posterior probability of the initially learned (blue) context declines, while that of the novel context (gray line in Fig 3B) associated with uniform contingencies *ϕ*^novel^ increases. After 3 trials, the novel context becomes the most likely explanation with a posterior probability *q*(*c*_*K*+1_ ≥ 0.5) and in response the agent opens a new context, now believing there are *K* = 2 contexts in the environment. It expands its repertoire of memorized reward contingencies by adding new contingencies *ϕ*_2_ associated with context 2. As the agent now believes with high certainty that it is in this new (orange) context, it can quickly learn these contingencies without having to forget or overwrite all the information learned for context 1 (Fig 3A bottom). Correspondingly, the agent begins to consistently perform the correct action of choosing bandit 2 in context 2 within 5 trials after the true latent cause switch (not shown).

The agent can also successfully identify when a previously learned context reoccurs, rather than further opening new contexts, as shown from trial *τ* = 200 onward in Fig 3B. In trial *τ* = 200, the agent encounters the previously experienced context 1, where bandit 1 elicits the rewards with a high probability again. In response, surprise increases for context 2, decreases for context 1 and briefly increases before declining again for the novel context. Correspondingly, the agent correctly infers that the most likely explanation is the reoccurrence of context 1 and assigns a high posterior probability to this context. Similarly, in trial *τ* = 300 the contingencies switch again to those of context 2, which the agent correctly identifies.

This example illustrates how in principle the NP-BCC utilizes contextual structure learning to create a growing repertoire of behaviourally relevant contextual memories via surprise minimisation. It also shows how the contextual inference principle guides structure learning, with contextual uncertainty estimates being used to both probe whether something novel has taken place in the environment, and to update associated contingencies such as the probability of receiving a reward in a given state and context *p*(*r*|*s, c, ϕ*).

### Contextual inference principle and flexibility vs stability

Beyond the singular example above, additional simulations show that an NP-BCC agent can reliably acquire the contextual structure of an *M* = 2 and *M* = 3 MAB task, showing that the proposed inference scheme is robust. See Fig 4 for the inferred posterior over contexts for 2- and 3-armed MAB tasks, averaged over agent instances.

**Fig 4.**
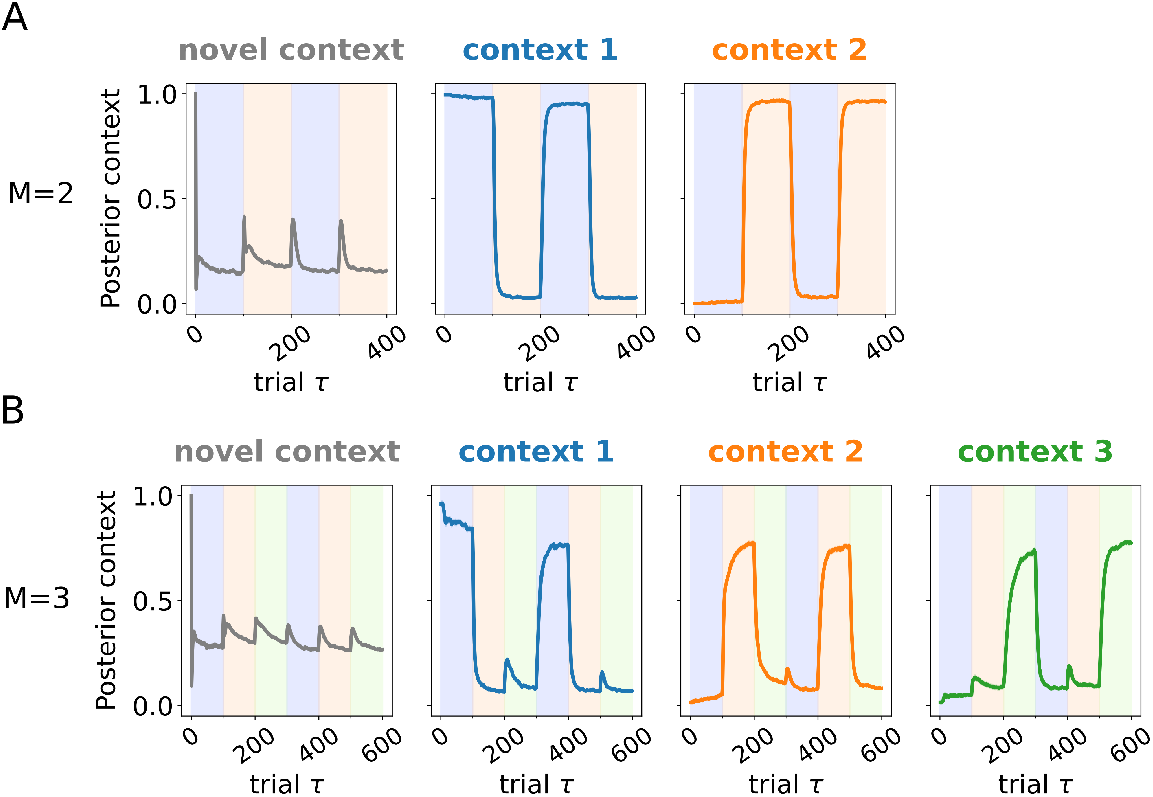
Characteristic contextual inference dynamics in *M* = 2 and *M* = 3 MAB tasks. **A)** The panel depicts the inferred novel context probability *q*(*c*_*τ*_ = *K* + 1) in gray and the renormalised posterior probability *q*^*′*^ (*c*_*τ*_) of each context in colour (as in Fig 3) across trials in an *M* = 2 MAB task, averaged over 200 agent instances. Coloured shaded areas indicate trials at which context 1 or 2 was active. Agents correctly learned the contextual representation of the task, as indicated by the increasing posterior during trials when that context is active. **B)** Same as in A but for a *M* = 3 MAB task. Colour coding is the same as above where the posterior over the third context is plotted in green. Agents again correctly inferred the number of active contexts and when they were active.

Averaging across agents illustrates how behavioural stability control is reflected in the dynamics of contextual inference. As indicated in the previous section, contextual inference reactivates previously learned contextual memories when they become relevant again. Because the NP-BCC weights the expression of contextual memories by their posterior probability, behaviour becomes more stable when a single context dominates the posterior and more flexible when several contexts remain plausible for a short period following a context switch in the task environment.

These characteristic contextual inference dynamics are visible at and around context change points in Fig 4. Each time a context switch occurs in the task environment, agents experience heightened surprise. This leads to an increase in contextual uncertainty and a switch away from a behavioural stability mode. For example in Fig 4B after the context switch at *τ* = 400, the posterior probability of the previously active context context 1 decreases and that of context 2, context 3 and the novel context start rising. This low contextual certainty state leads to a transient probing of different contextual memories to gather evidence for possible contexts. With sufficient probing the posterior probability of the inactive contexts drops again (context 3) and that of the context that reoccurred (context 2) sharply rises. This increase in contextual certainty corresponds to a transition back to a stability mode, where agents primarily execute actions optimal under the currently inferred active context and allocate belief updates about contingencies to that context, rather than further probing existing memories.

Additionally, similar to animals and humans [9, 22, 59–61], NP-BCC agents gain certainty about the currently active context faster when a context has been encountered before as compared to when it is encountered for the first time. To quantify this, in Fig 5 we compare the trial at which agents first attain a posterior probability *q* (*c* = *c*_true_) *>* 0.75 for the currently active context, depending on whether the context was newly encountered or not. We found that, on average, agents inferred the occurrence of a context significantly faster relative to the initial encounter (Mann–Whitney–Wilcoxon test, two-sided: *U* = 1.09 *×* 10^5^, *p* = 1.86 *×* 10^−18^).

**Fig 5.**
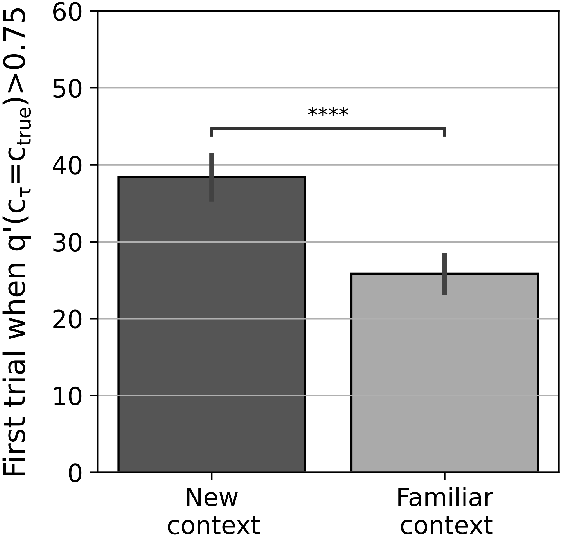
Recognition speed for new versus familiar contexts. The figure shows the trial at which agents achieved high context certainty (*q*^*′*^ (*c*_*τ*_ = *c*_true_) *>* 0.75) for the currently active context, depending on whether the active context was previously encountered or novel. A Mann–Whitney–Wilcoxon test indicated that familiar contexts were inferred significantly faster than novel ones (*U* = 1.09 × 10^5^, *p* = 1.86 × 10^−18^). Agents used for analysis were the same agents performing an *M* = 3 MAB task in Fig 4B. Data for encountering context 1 was not used, since it is immediately inferred in the very first trial (see Methods, Section Inference procedure).

Lastly, 99% of 200 agents successfully learned the contextual representation of the *M* = 2 task whereas only 82% of 200 agents learned the more complex *M* = 3 task for the same amount of training. Agents were considered to have successfully acquired the contextual representation of a task if the true currently active context *c*_true_ had the highest posterior probability on more than 90% of trials following the structure acquisition phase of a simulation (after *τ* = 200 in Fig 4A and *τ* = 300 in Fig 4B). Agents performing the *M* = 3 task also achieved overall lower posterior certainty for the currently active context compared to the *M* = 2 task. Increasing the amount of training to 130 trials per context in the *M* = 3 task brings the proportion of agents that successfully acquired the task structure back up to 97% (data not shown). Even though overall agents can learn the correct contextual structure of the *M* = 3 task, these simulations already hint at the increased data requirements in more complex environments for naive structure learning agents, shown in detail in the next section.

### Limits of naive rational agents

As demonstrated above, agents so far learned contextual representations in an *M* = 2 and *M* = 3 MAB task and used them to flexibly switch between task demands. However, we found that this naive structure learning starts to err on the side of flexibility with increasing task complexity, with agents remaining overly uncertain about the active context, which can lead to aberrant context opening, erroneous structure learning and decreased behavioural performance, as observed for 18% of agents in the 3-armed MAB tasks. When we further increased task complexity to *M* = 4 choice options (Fig 6A), pure structure learning becomes insufficient for acquiring the contextual task structure for the same amount of training. This can be seen by the rather low posterior over contexts, often around only 0.5 for context 3 and 4. We found that agents learn the contingencies associated with each context when extending the training to 300 trials per context in the *M* = 4 MAB task (Fig 6B) compared to a 100 and 130 trials per context in the *M* = 2 or *M* = 3 tasks (Fig 4B).

**Fig 6.**
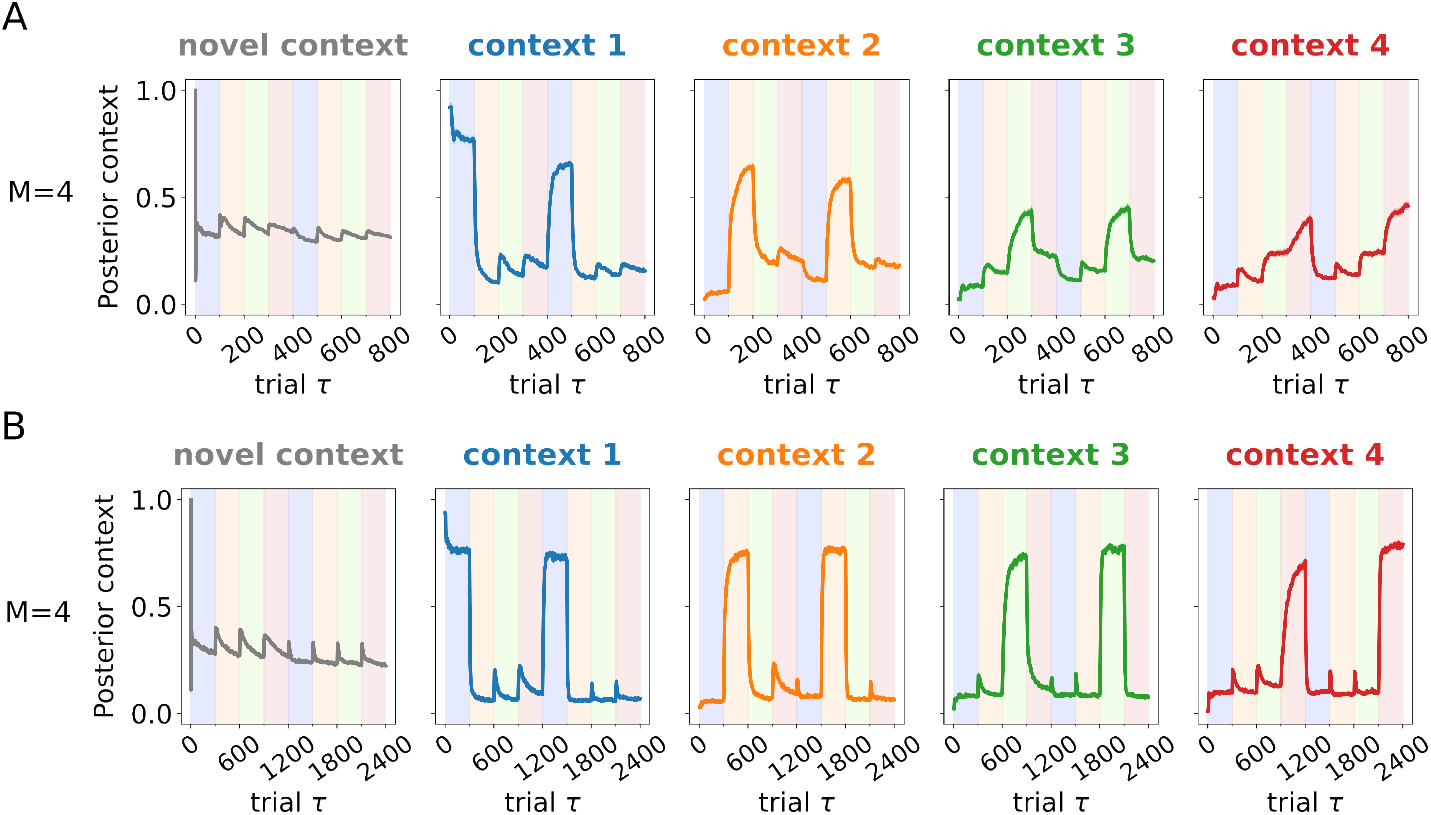
Naive structure learners require longer training times in more complex environments. **A)** Plots show the inferred novel context probability *q*(*c*_*τ*_ = *K* + 1) in gray and the renormalised posterior over contexts *q*^*′*^ (*c*_*τ*_) in colour (as in previous figures) when performing an *M* = 4 MAB task with a shorter training regime, averaged over 200 agent instances. Here each context is active for a 100 trials and all contexts are encountered twice. Colour coding for the different contexts is the same as in previous figures, with the extra fourth context plotted in red. Shaded regions indicate the currently active context. In this short training regime, context inference became unstable and agents are unable to correctly learn the contextual representation of the task, having only properly learned the contingencies associated with context 1. Context mixing is evident, with contingencies associated with context 3 and 4 being partially learned into context 2 or into each other. This effect can be seen by the high posterior allocated to context 2 in trials when context 3 or 4 is active for example. **B)** Same as in A but with an extended training regime where each context lasted 300 trials. Extending the exposure from 100 to 300 trials per context substantially reduces the amount of context mixing. Agents are capable of inferring the correct number of active contexts and when they became active.

Higher training requirements arise because once agents learn the optimal response in a context, they exploit this knowledge and rarely probe suboptimal bandits. As a consequence, the contingencies associated with suboptimal bandits remain under-learned because learning signals are only generated when a bandit is selected. Thus, although the agent has learned that an action is suboptimal, the corresponding contingencies are updated far less frequently than those of the optimal action. Then, when a context switch occurs in the environment, selecting under-learned bandits generates a relatively lower surprise under the previously active context. This can lead to erroneously learning contingencies of the newly active context into previously active ones, which in turn leads to increased contextual uncertainty, because agents have acquired the wrong task structure. Such context mixing can be observed in Fig 6A, where the probability for context 2 (orange line) increases during trials when context 3 or 4 is active in the environment, indicating that the contingencies for these contexts were partially learned into context 2.

Motivated by these limitations of naive structure learning, we next examined whether repetition-based automatisation and utilizing structured prior knowledge could support more robust performance in familiar tasks and more rapid acquisition of novel task structure. The general idea is that repetition-based automatisation may lend more stability and more certainty to contextual inference, while prior knowledge may help with rapid acquisition of novel contexts. Given our observations in the 4-armed MAB task, these potential improvements may increase the overall performance of the naive NP-BCC agent. We therefore implemented these two cognitive mechanisms in the NP-BCC model as a context-specific prior over policies and template contexts to test whether they can rescue performance in more complex environments like the *M* = 4 MAB task where naive agents fail.

### Automatisation stabilizes contextual inference

Here we show that automatisation has a significant stabilizing effect on structure learning and context certainty in the *M* = 4 MAB task, for the long (300 trials) training regime. In the previous sections, behaviour was chosen primarily based on value-based information in the form of reward contingency-based surprise signals derived from the context- and policy-specific free energy (Eq (9)). Now, in addition, the NP-BCC agent has the capability to learn context-specific automatisms in the form of a context-dependent prior over behaviour (Eq (8)). This prior assigns an a priori bias to repeat policies in the same context and is proportional to the number of times the policy has been performed in the past [14]. How fast the prior changes in response to repetition is governed by a parameter *h* ∈ (0, 1), where higher values of *h* result in a larger bias with fewer repetitions (see Methods, Section Automatisation learning and automatisation tendency). Actions are chosen from the posterior over policies which is obtained by combining this prior with a likelihood containing reward information from the policy-specific free energy that favours policies that are likely to lead to desirable outcomes. Note that in the MAB task, policies map directly onto actions, since only a single choice is made per behavioural episode. In summary, the NP-BCC formalizes automatisation as repetition-based learning which is integrated into value-based processing via Bayesian belief update.

In Fig 7A we compare the contextual inference traces of non-automatising agents with an *h* ≈ 0 (in black) to those of moderately automatising agents with an *h* = 0.025 (in colour) performing the same *M* = 4 MAB task as in Fig 6B. Note that the *h* ≈ 0 agents are the same as those shown in Fig 6. Automatisation enabled learning low uncertainty contextual representations faster, as indicated by the more rapidly declining posterior probability for the novel context after each context switch. Additionally, moderately automatizing agents achieved higher contextual certainty overall compared to naive agents, as indicated by the higher posterior probability allocated to the most likely context. This effect emerges because learned behavioural biases are a part of an agent’s contextual representation (Fig 1). Concretely, the executed policy becomes informative for what context agents believe they are in. This is because in automatising agents the prior over policies varies with the specific context and reflects learned biases. The executed policy therefore serves as an additional source of surprise and contributes to context inference and context certainty via the policy cross-entropy term in Eq 11. In non-automatising agents, the context-specific prior over policies remains flat, hence this term does not differ between contexts and does not contribute to context inference.

**Fig 7.**
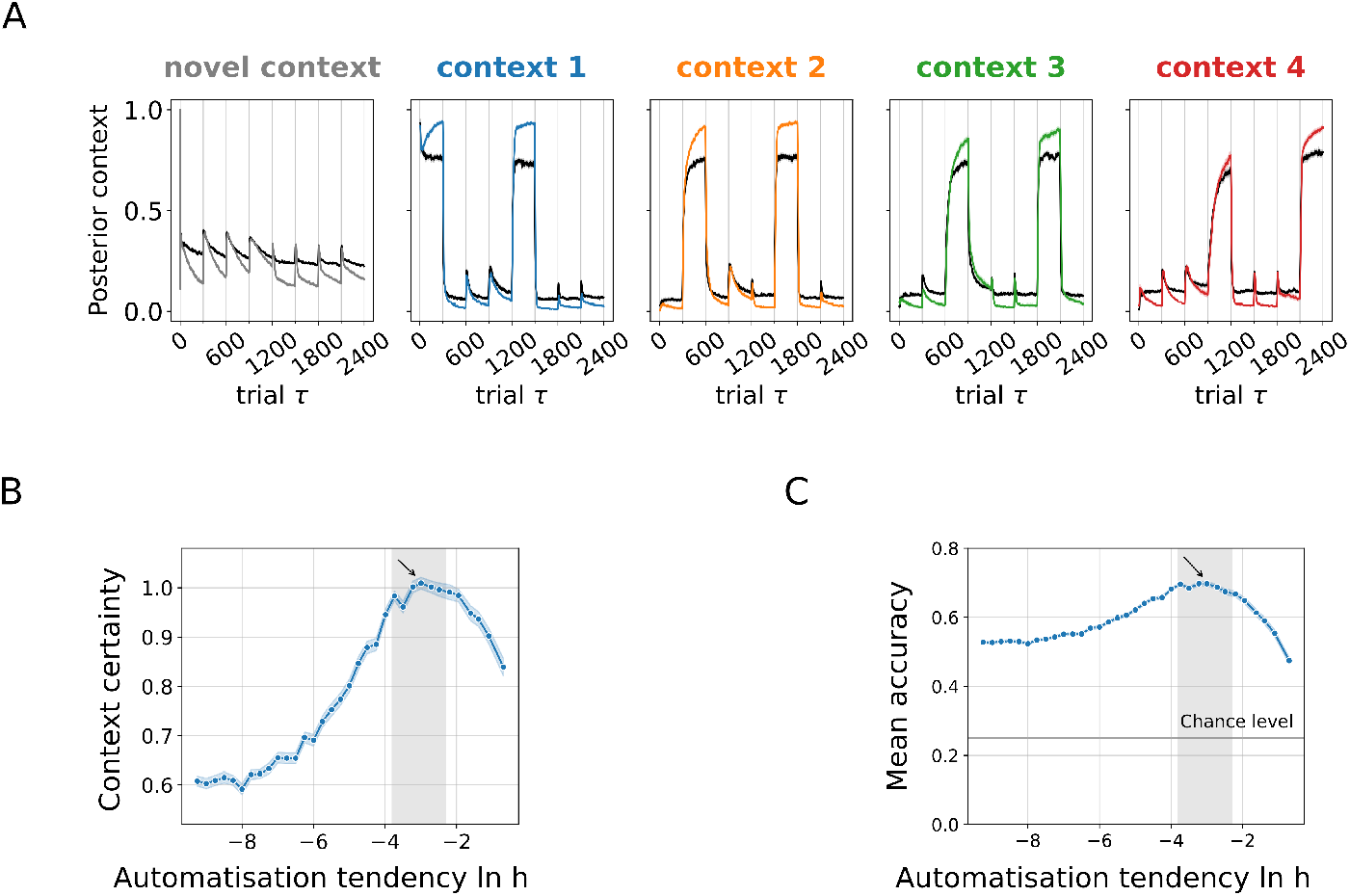
Automatisation stabilizes structure learning and contextual inference. **A)** The panel compares the inferred posterior distributions over contexts for non-automatising agents (shown in black; automatisation tendency *h* = 0.0001) and moderately automatising agents (context colour coding same as in other figures, *h* = 0.025). Moderately automatising had a more rapidly decreasing posterior probability for the novel context after a context switch and inferred the currently active context with a higher certainty compared to non-automatising agents. Results indicate that automatising agents infer contextual representations more rapidly and with higher certainty. The data for non-automatising agents was taken from Fig 6. Data is averaged over 200 agent instances. **B)** The panel shows the context certainty (formalized as the relative entropy *D*_*KL*_[*q*(*c*_*τ*_)|*p*_unif_(*c*_*τ*_)] between the inferred posterior over contexts *q*(*c*_*τ*_) and a uniform distribution over possible contexts *p*_unif_ (*c*_*τ*_)) as a function of automatisation tendency *h* plotted in log space. Values were averaged for trials after the initial structure acquisition (*τ >* 1200) and 200 agent instances. Moderately automatising agents with ln *h* between −3.8 and −2.3 (*h* between approximately 0.022 and 0.1) achieve the highest context certainty overall. The arrow indicates the automatisation strength *h* = 0.025 used for moderate automatisation tendency agents in the plot above. **C** The panel shows mean accuracy, measured as the proportion of trials in which the agent executed the optimal policy for the active context, as a function of log automatisation tendency ln *h*. Values were again averaged for trials after the initial structure acquisition (*τ >* 1200) and 200 agent instances. The arrow indicates the automatisation strength *h* = 0.025 used for high automatisation tendency agents in the **A**.

Correspondingly, agents generally achieve higher context certainty when they automatise behaviour in stable contexts, because the execution of the automatised policy itself adds more evidence for the associated context. Hence automatisation not only aides the exploitation of known contexts but also supports contextual structure learning and inference by virtue of being part of the contextual representation of the world.

This effect can be clearly seen in Fig 7 which shows how different degrees of context-specific behavioural automatisation can be more or less beneficial for contextual inference in terms of context certainty (Fig 7B) and behavioural optimisation in terms of accuracy (Fig 7C). Systematically varying the automatisation tendency in the NP-BCC yielded U-shaped relationships between automatisation and both contextual certainty and behavioural accuracy, in which an optimal degree of automatisation provided behavioural stability in complex environments while preserving flexibility under changing conditions.

Agents with a moderate degree of automatisation (shown in shaded figure range), at which the surprise generated from the executed policy is pronounced but does not outweigh other surprise signals, achieve the highest overall context certainty (Fig 7B) and the highest accuracy (Fig 7C). Conversely, agents with negligible automatisation which corresponds to highly negative values of ln *h* (e.g. ln *h <* −8) exhibit lower contextual certainty (Fig 7B) and correspondingly less well-defined contextual representations. The accuracy of these agents is still high compared to chance level but lower than that of the moderately automatising agents which benefit from the higher contextual certainty and learned behavioural preferences (Fig 7C).

At the other extreme, when agent’s automatisation tendency is too high, they develop an excessive behavioural bias, which prevents them from inferring a context switch and leads to suboptimal behaviour. Such agents with *h* approaching 1 (and ln *h* approaching 0) have a relatively high certainty in what context they are in compared to non-automatising agents (Fig 7B) because they believe they are in the context where their preferred policy is optimal, whatever the true context is. Naturally, their accuracy however is low (Fig 7C), because they are prone to assuming the incorrect contextual contingencies. This illustrates that high contextual certainty can coexist with an incorrect contextual representation. Accordingly, having a higher context certainty does not necessarily imply learning the correct contextual representation of a task. Interestingly, this is what is observed in the habit literature as well with overly habitual animals unable to infer uncued context switches [61–63].

Taken together, these results show that repetition-based automatisation can stabilize behaviour under increasing task complexity. However, automatisation alone does not substantially reduce the amount of experience required to acquire accurate contextual representations in more complex environments, such as the *M* = 4 MAB task. This suggests that additional mechanisms may help to support faster contextual structure learning.

### Template contexts aide novel context acquisition

In the analyses above, agents detected the presence of a new context by tracking the surprise generated by a novel context associated with uniform contingencies. The novel context contingencies *ϕ*^novel^ (Fig 3A) exemplify the belief that a new context, about which the agent knows nothing, has occurred. Once the agent decides something novel happened, it instantiates a new context and learns its associated contingencies in the following trials. This initial learning is an iterative, potentially slow process which naturally takes longer in more complex environments, as, for instance, in the *M* = 4 MAB task (Fig 6B).

However, humans are able to adapt to and learn the contingencies of a new environment within a few encounters, which is qualitatively different to a slow learning process. To achieve such a rapid acquisition, humans use previous knowledge about what actions might be useful in a new situation, which in cognitive psychology is often referred to as schemata learning and deployment [24–26, 33]. In order to enable the NP-BCC agents to emulate this ability, we use template contexts which allow agents to populate a novel context with prior assumptions about its structure.

For demonstration purposes, we adopt a simple Occam’s razor-based prior [64, 65] for the MAB tasks that one bandit is rewarding, while all others are not, which will act as the agent’s first guess how the environment is structured, and will then be successively updated with each experience of the environment. We therefore introduce a set of initial template contingencies 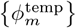 which imply a high reward probability for bandit *m*, and low probabilities for all other bandits (see Methods, Section Template contexts and reward hyperparameter initialization). NP-BCC agents which use template contexts detect the presence of a novel context in the same way as naive structure learners, i.e., by tracking the probability of a novel context defined by uniform contingencies *ϕ*^novel^ (Fig 3A). The difference lies in how template-using agents open a new context: its contingencies *ϕ*_*K*+1_ are initialized by selecting the template contingencies 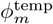 that yield the lowest surprise given the observation responsible for the context’s creation. To not conflate effects caused by use of templates with those due to automatisation, we used non-automatising agents (*h* ≈ 0) in the following simulations.

We found that agents that incorporated prior assumptions via template contexts acquired the contextual structure of an *M* = 4 MAB task with extended training faster and more reliably compared to naive structure learners. Concretely, agents inferred to be in a newly encountered context with a probability *q*(*c* = *c*_true_) *>* 0.75 substantially faster (Fig 8A). As expected, template contexts primarily aided initial contextual structure acquisition and conferred smaller benefits when encountering an already learned context (Fig 8B). Additionally, 99% of simulated agents that utilized template contexts correctly learned the task structure, compared to 94% of naive structure learning agents.

**Fig 8.**
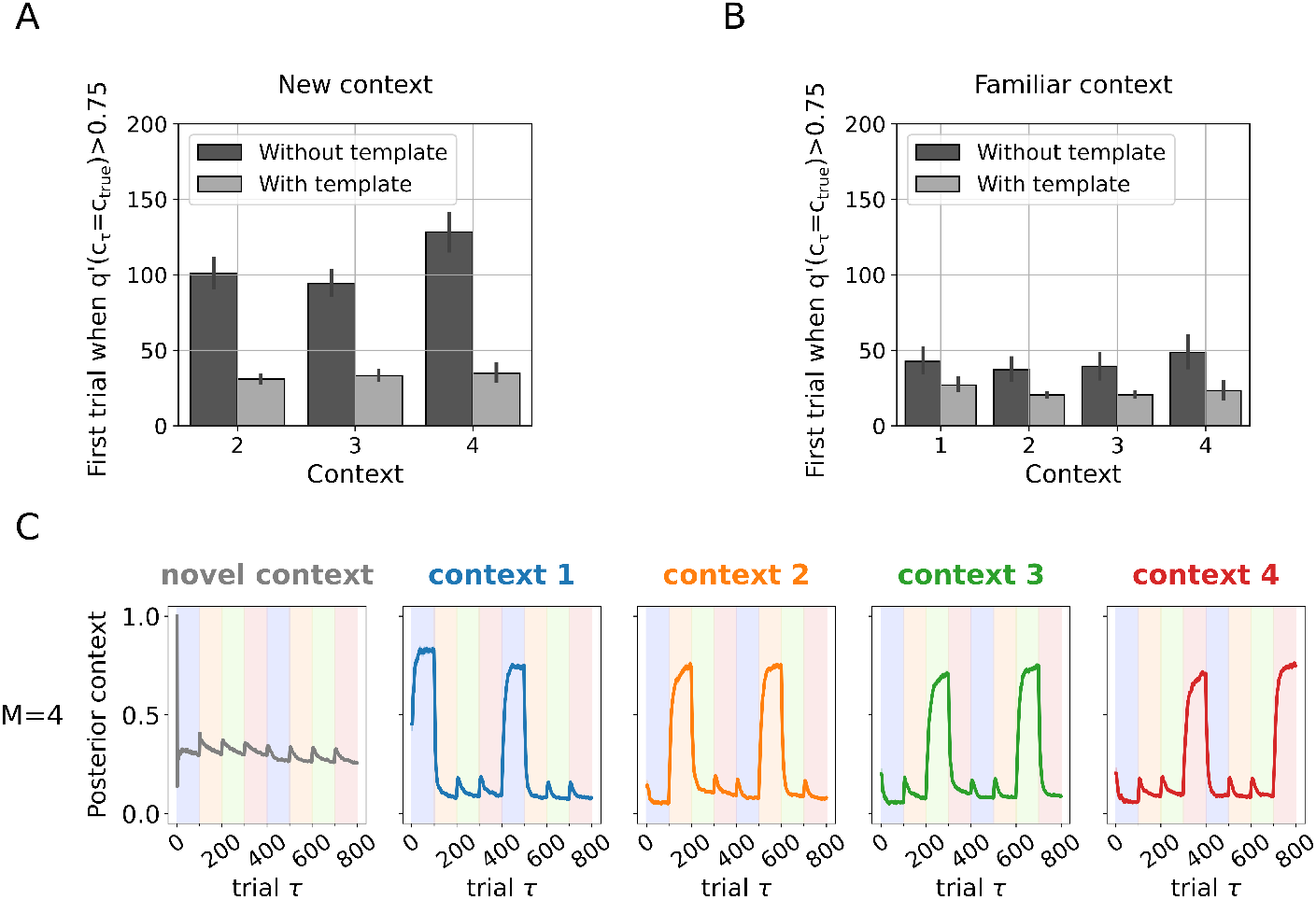
Template contexts speed up and stabilize contextual inference. **A)** Plot shows data from 200 agents (with or without template contexts) performing a 4-armed MAB task with extended training as in Fig 6. The currently active context is shown on the x-axis and the first trial at which agents inferred the correct context with a renormalised probability *q*^*′*^(*c*_*τ*_ = *c*_true_) *>* 0.75 on the y-axis. Only trials when agents encounter a given context for the first time are used. The plot compares agents performing naive structure learning without templates contexts (shown in dark gray) to agents using templates contexts (shown in light gray). Template-using agents inferred the presence of a new context with high certainty within about 25 instead of about 100 trials. Data for agents that do not use template contexts was taken from agents shown in Fig 6B. Both types of agents did not use automatisation (*h* = 0.0001). Because agents are initialized to start with an empty set of known contexts and open one at the first trial, data for context 1 was not used in the analysis, since it is not informative. **B)** This plot compares, for previously encountered, i.e., familiar, contexts, at which trial the two types of agents infer the correct context with a renormalised posterior probability *q*^*′*^(*c*_*τ*_ = *c*_true_) *>* 0.75 for the first time. Template contexts also improve speed of context recognition upon repeated context presentation. This benefit is less, compared to when the context is newly encountered. **C)** Plot shows the renormalised posterior over contexts in a *M* = 4 MAB task, averaged over 200 agent instances. Agents deployed context templates but did not use automatisation (*h* = 0.0001). Colour coding and meaning of vertical lines is the same as in previous figures. Deployment of template contexts stabilized context inference and prevented context mixing compared to the naive structure learning case shown in Fig 6B, as evidenced by the fact that all 200 agents inferred the correct number of contexts with high certainty and successfully inferred switches between them at the correct points in the task.

Importantly, template context deployment enabled agents to acquire the contextual structure of an *M* = 4 MAB task with a short training regime of 100 trials per context (Fig 8C), indicating their usefulness in rapidly acquiring novel tasks.

Clearly, it is an open question how exactly these template contexts would be learned or deployed sensibly for tasks different from the current MAB task (see Discussion). Here, we hard-coded appropriate prior knowledge for a specific task, where other tasks may need different priors.

Nevertheless, these results provide a proof of principle and showcase how incorporating prior knowledge can aid decision making in complex dynamic environments where previous experience suggests plausible candidate contextual structures. The addition of structured assumptions into newly opened contexts helps restore the stability-flexibility balance even in complex environments where naive rational agents require substantially longer training times to acquire task structure and high contextual certainty.

## Discussion

In this work we present a computational model of contextual inference for value-based decision making that integrates nonparametric structure learning with mechanisms that formalize a behavioural stability and flexibility trade-off. Through simulations, we illustrate how the resulting nonparametric Bayesian Contextual Control (NP-BCC) model can acquire and reuse an expanding repertoire of latent contexts and their transition dynamics through surprise-driven inference, which in turn supports flexible adaptation to changing task demands while preserving previously learned task representations. These results are intended as mechanistic proof-of-principle demonstrations of how such interacting processes can be formalized computationally.

Our simulation results indicate that contextual inference and structure learning in the NP-BCC model can mediate the trade-off between behavioural flexibility and stability by regulating when stored contextual memories are expressed, updated, or newly formed, consistent with previous contextual inference models [9, 13, 22]. Importantly, we further show that incorporating well-established cognitive mechanisms known to underlie human decision-making, specifically repetition-based automatisation and schema-like structured prior knowledge, stabilizes contextual inference and structure learning in more complex environments. These mechanisms reduce erroneous context formation, improve performance, and markedly decrease the data required to acquire accurate task representations, stressing their functional role in supporting adaptive behaviour under increasing task complexity.

### Integrating automatisation and schema-like representations within a contextual inference framework

Automatisation is traditionally formalised as a context-dependent stimulus-response association [30, 66]. Computationally it is viewed as an efficient heuristic that can optimize behaviour in stable environments but also lead to maladaptive behaviour in dynamic environments when expressed excessively. Our results broadly support this view and further clarify the functional role automatisation may play in adaptive behaviour. Contextualizing automatisation did not simply help optimize behaviour in stable contexts (Fig 7). In the NP-BCC automatisation is learned in a context-specific manner, such that different contexts become associated with distinct behavioural biases. Crucially, as a result, action execution itself becomes informative about the currently active context: executing an action that is strongly expected under a given context provides evidence in favour of that context, analogous to how observed rewards inform contextual beliefs. This additional, self-generated source of evidence contributes both to stabilising context inference and to acquiring an appropriate contextual structure during learning.

To further unpack this mechanism, it is useful to consider how contextual representations are traditionally understood in learning theory, where contextual interpretations are typically assigned to stimuli rather than actions. For example in reacquisition, a previously extinguished response is relearned more rapidly when the conditioned stimulus and unconditioned stimulus are paired again [67, 68]. In reinstatement, an extinguished response can return following presentation of the unconditioned stimulus alone [69, 70]. Both phenomena are commonly understood as a consequences of implicitly defined contextual representations, in which the presentation of associated stimuli serves as a contextual cue that signals the original acquisition context [10]. Similar to its parent model [14], the NP-BCC extends this inferential logic to actions. When actions themselves become part of the implicit contextual representation (via repetition-based, context-specific action priors), their execution serves as an internal cue that signals the associated context. The NP-BCC formalizes this bidirectional flow of information, in which context informs behaviour and behaviour informs possible contexts while mechanistically modelling the process of acquiring a contextual representation in the first place.

In addition to automatisation, the NP-BCC explores how schema-like structured prior knowledge can be embedded into newly opened contexts via template contexts. In the model, template contexts encode abstract, behaviourally relevant regularities such as the presence of a unique optimal choice, rather than specific stimulus-response contingencies. This allows newly inferred contexts to inherit structured expectations about task organization, thereby supporting rapid learning and stabilization of behaviour in complex environments.

From a cognitive perspective, these template contexts closely resemble what are commonly referred to as schemas: abstract knowledge structures acquired over multiple experiences that organize prior knowledge and guide perception, learning and action [26, 33, 71]. Schemas are typically characterized by their associative and abstract nature, their basis in multiple episodes and their lack of single-instance detail. In this sense, template contexts can be understood as learned, generalized representations that capture relationships between task variables such as states and rewards. Consistent with theoretical accounts suggesting that schemas support efficient reinforcement learning in dynamic environments [72], our results indicate that embedding such structured priors facilitates both context acquisition and performance stabilization.

Although the NP-BCC is a computational model, its unification of contextual inference and schema-like representations naturally raises the question of whether these processes also overlap at the neural level. There is a lot of evidence relating the at first glance disparate fields of contextual inference and schemas [33, 71]. Even though there is no broadly agreed upon circuit level explanation of contextual inference and schema formation and deployment, there appears to be significant neuroscientific evidence for overlapping neural substrates [9, 71]. For example, areas such as the medial prefrontal cortex (mPFC), specifically the orbitofrontal cortex (OFC) and ventromedial prefrontal cortex (vmPFC) appear to be involved both in contextual inference [15, 22, 73–76] and schema instantiation and deployment [77–79], supported by broader interactions with the hippocampus and posterior cortical regions [71, 77, 80, 81]. There are competing theories regarding the functions of these regions. Although the OFC and vmPFC have often been described in terms of value representation [82, 83], more recent theoretical and empirical work argues that these regions encode rich task representations that go beyond simple value signals, consistent with the hypothesis that prefrontal OFC and vmPFC support latent task states and hidden variables that underlie contextual inference and adaptive control [73, 84–86].

Correspondingly, it has recently been proposed that the same principles that support flexible and adaptive instrumental behaviour, such as prediction error-driven learning, hierarchical representation acquisition and latent state inference, also govern the learning of schemas [33]. The NP-BCC instantiates many of these principles by treating contextual inference over latent states within a hierarchical representation, guided by surprise minimisation. This formulation provides a quantitative framework for studying schema formation and deployment and how these processes interact with contextual inference and instrumental learning.

### Implications for mental disorders such as substance use disorders

Aberrant contextual processing has been proposed to underlie a variety of clinical conditions such as schizophrenia, post-traumatic stress disorder and substance use disorders (SUDs) [17]. Substance use disorders for example have been widely associated with maladaptive persistence of context-dependent behaviour, where learned substance use patterns remain strongly coupled to consumption-related contexts despite adverse consequences [87]. Our results suggest potential computational mechanisms through which such persistence may arise. In the NP-BCC framework, repetition-based automatisation strengthens context-specific action priors. Hence, repeatedly executed substance use behaviours both stabilize performance and provide additional evidence in favour of the currently inferred context. While adaptive under stable conditions, excessive coupling between action repetition and contextual belief updating could contribute to self-reinforcing inference loops in which habitual behaviour biases context inference toward previously learned consumption contexts. This can in turn increase susceptibility to craving and relapse [88].

In parallel, schema-like template contexts illustrate how structured prior expectations can accelerate context acquisition and stabilization. Although beneficial for efficient learning, strong prior assumptions can also bias how novel situations are interpreted [71]. Hence, the deployment of schematic knowledge is another potential computational mechanisms that could favour the reinstatement of familiar maladaptive contextual structures linked to craving or relapse.

Seen through this lens, substance use disorders could be understood as a pathological shift in the stability–flexibility dynamics formalized by the NP-BCC, where contextual inference becomes biased toward rigid persistence rather than adaptive updating. Importantly, this interpretation is not intended as a clinical model, but as a formal account that highlights candidate mechanisms linking contextual inference, repetition-based automatization, and structured priors to behavioural rigidity. Future work could use this framework to experimentally probe how these mechanisms shape trajectories of losing and regaining control in SUDs, providing a formal basis for testing mechanistic hypotheses about craving, relapse, and longer-term behavioural dynamics [89, 90].

### Relation to computational models of structure learning

Previous nonparametric contextual models have initialized novel contexts in various ways. For example, the COIN motor learning model [9] employs a particle filtering approach to approximate the posterior over latent variables and model parameters, allowing each particle to independently instantiate new contexts at any trial. When a new context is created, its dynamical parameters are sampled from corresponding hyperpriors, and the associated state is initialized to the steady state of the resulting linear dynamical system.

In the value-based decision making model by [22], the authors took a different approach. The model tracked a limited set of contexts in a working-memory buffer and initialized a novel context when all monitored contexts showed low reliability, i.e., had poor explanatory power. A new “probe” context was then generated by combining contexts from long-term memory with added noise and was subsequently evaluated for reliability. In the NP-BCC, this idea is extended by explicitly coupling the creation of novel contexts to the deployment of schema-like priors, allowing newly opened contexts to inherit structured expectations about task organization.

Recent Active Inference models of structure learning have explored mechanisms for bounded state space expansion and reduction within a single generative model [91]. However, these approaches did not incorporate an explicit contextual layer or nonparametric priors over latent structure. In contrast, the NP-BCC integrates an Active Inference based approach with a nonparametric prior over contexts, allowing flexible inference over an unbounded contextual structure. While the NP-BCC is grounded in an Active Inference framework and is therefore in principle capable of planning, exploring planning behaviour was beyond the scope of the present study and remains an important direction for future work.

### Limitations and future directions

In the present implementation of the NP-BCC, template contexts were specified a priori as a minimal instantiation of schema-like structured prior knowledge. Once instantiated, templates were consolidated immediately, allowing us to isolate and examine how structured priors interact with contextual inference and structure learning. This simplified treatment was intentionally chosen to establish a clear conceptual link between schema representations and contextual inference within a unified computational framework. Importantly, schema formation, selection, and deployment are likely to involve richer dynamics than captured here, including the evaluation of candidate schemas and adaptive loading and unloading based on task demands, as suggested by prior work [22]. As briefly mentioned above, recent theoretical and empirical work suggests that schema formation and deployment are also likely driven by processes such as prediction error–based learning and latent state inference, in concert with dimensionality reduction and state abstraction [33]. Elaborating these dynamics and formalizing their relationship to surprise minimization constitute important directions for future research.

An important aspect of the NP-BCC concerns how model stability depends on a small number of hyperparameters that regulate contextual structure learning. In particular, the parameters *γ, α*, and *κ* control the tendency to instantiate novel contexts through the global and local priors over contexts, as well as the strength of self-transition bias, respectively. Together, these parameters determine the agent’s prior expectations about the number of latent contexts and the degree of environmental stability. In practice, identifying parameter regimes that supported stable online contextual structure learning within a variational inference framework required task-specific adjustments (see Table S1 in Supplementary material). This sensitivity is expected and reflects the fact that these hyperparameters encode strong a priori assumptions about task complexity and environmental volatility, which differ across environments and directly shape whether the agent favours context creation, reuse, or persistence. In biological systems, such strong assumptions are likely regulated by higher-level adaptive processes operating over developmental or even evolutionary timescales. From this perspective, the observed sensitivity points to the absence of a higher-level adaptive mechanism that regulates these priors, and would be a natural direction for future model extensions. Nonetheless, we expect the main findings of this paper to generalize to other tasks, both with single and sequential choices, as long as the hyperparameters are chosen in accordance with the task environment, because the main effects are driven not by the hyperparameters but by the mechanisms encoded in the NP-BCC.

Hyperparameter values also had a strong effect on the dynamics of contextual inference, as they regulate how quickly the agent gains certainty in the contextual structure of the environment and its transition dynamics. Misspecifying these values could lead to persistently lower contextual certainty even when the correct structure was ultimately acquired. This sensitivity reflects how strongly prior assumptions about environmental stability and complexity shape the agent’s ability to commit to and maintain contextual beliefs.

Another potential limitation of the current implementation concerns its reliance on strictly online belief updating. How humans learn the transition dynamics of an environment remains an open question, but there is substantial evidence that memory reactivation during sleep supports consolidation and the integration of new information into existing schemas [92, 93]. This is consistent with the possibility that aspects of contextual structure learning rely on offline, batch-like updating rather than exclusively on moment-by-moment inference, as implemented in the NP-BCC. From this perspective, trial-by-trial online Dirichlet count learning may provide an incomplete computational formalisation of how transition structure is acquired, motivating future work that incorporates offline or retrospective updating mechanisms.

A related limitation concerns the robustness of the context opening mechanism under approximate online inference. As illustrated in Results, the NP-BCC can become unstable when novel contexts are instantiated prematurely or when uncertainty about a potential context switch persists without resolving decisively. In such cases, observations from a newly emerging context may be incorporated into an incorrectly inferred active context, leading to transient or sustained misassignment. Importantly, our results demonstrate that this form of instability is substantially reduced when agents employ template contexts, indicating that structured priors play a stabilizing role during contextual transitions. More generally, these findings suggest that robust context creation likely requires integrating evidence over multiple observations and coupling context opening more tightly to schema loading and formation processes. Mechanistically elaborating these components in a more principled manner, as well as extending the current approximation toward a full HDP-HMM prior, may further improve robustness.

## Conclusion

In summary, we introduced the nonparametric Bayesian Contextual Control (NP-BCC) model as a computational framework for contextual structure learning in value-based decision making. The model integrates repetition-based automatisation and schema-like structured priors into a common account of contextual inference. Our results indicate a fundamental role for automatisation when it is formalised as part of a latent contextual representation, demonstrating how context-specific action biases acquired through repetition can contribute not only to efficient action selection but also to the stabilisation of contextual beliefs. More broadly, the NP-BCC provides a principled link between contextual structure learning and schema formation, two literatures that have largely developed in parallel. By treating contexts as latent structures that can be inferred, reused, or newly instantiated, the model illustrates how agents can dynamically regulate the trade-off between behavioural flexibility and stability. In doing so, agents exploit established contextual representations when appropriate while remaining sensitive to evidence that calls for updating or restructuring task representations.

## Funding acknowledgements

This research was funded by the German Research Foundation (Deutsche Forschungsgemeinschaft https://www.dfg.de/, DFG project numbers 402 170 461 (TRR 265) - SK, MNS, SS; 454 245 598 (IRTG 2773) - MNS; and 521 379 614 (TRR 393) - MNS) and as part of Germany’s Excellence Strategy – EXC 2050/2 – Project ID 390696704 – Cluster of Excellence “Centre for Tactile Internet with Human-in-the-Loop” (CeTI) of Technische Universität Dresden - SK. The funders had no role in study design, data collection and analysis, decision to publish, or preparation of the manuscript.

## References

1. Goschke T, Bolte A. Emotional modulation of control dilemmas: The role of positive affect, reward, and dopamine in cognitive stability and flexibility. Neuropsychologia. 2014 9;62:403–23. doi:10.1016/j.neuropsychologia.2014.07.015.

2. French RM. Catastrophic forgetting in connectionist networks. Trends In Cognitive Sciences. 1999 4;3:128–35.

3. Musslick S, Bizyaeva A, Agaron S, Leonard N, Cohen JD. Stability-Flexibility Dilemma in Cognitive Control:A Dynamical System Perspective. Proceedings of the Annual Meeting of the Cognitive Science Society. 2019;41.

4. Mayr U, Grätz D. Does Cognitive Control have a General Stability/Flexibility Tradeoff Problem? Current Opinion in Behavioral Sciences. 2024;57:101389. doi:10.1016/j.cobeha.2024.101389.

5. Goschke T. Dysfunctions of decision-making and cognitive control as transdiagnostic mechanisms of mental disorders: Advances, gaps, and needs in current research. International Journal of Methods in Psychiatric Research. 2014 1;23:41–57. doi:10.1002/mpr.1410.

6. Giannone F, Ebrahimi C, Endrass T, Hansson AC, Schlagenhauf F, Sommer WH. Bad habits–good goals? Meta-analysis and translation of the habit construct to alcoholism. Translational Psychiatry. 2024 12;14. doi:10.1038/s41398-024-02965-1.

7. Ersche KD, Gillan CM, Jones SP, Williams GB, Ward LHE, Luijten M, et al. Carrots and sticks fail to change behavior in cocaine addiction. Science. 2016 6;352:1464–8. doi:10.1126/science.aaf0941.

8. Heinz A, Beck A, Halil MG, Pilhatsch M, Smolka MN, Liu S. Addiction as learned behavior patterns. Journal of Clinical Medicine. 2019 8;8. doi:10.3390/jcm8081086.

9. Heald JB, Lengyel M, Wolpert DM. Contextual inference underlies the learning of sensorimotor repertoires. Nature. 2021 12;600:489–93. doi:10.1038/s41586-021-04129-3.

10. Heald JB, Lengyel M, Wolpert DM. Contextual inference in learning and memory. Trends in Cognitive Sciences. 2023 1;27:43–64. doi:10.1016/j.tics.2022.10.004.

11. Heald JB, Wolpert DM, Lengyel M. The Computational and Neural Bases of Context-Dependent Learning. Annual Review of Neuroscience. 2023 7;46:233–58. doi:10.1146/ANNUREV-NEURO-092322-100402.

12. Butz MV, Mittenbühler M, Schwöbel S, Achimova A, Gumbsch C, Otte S, et al. Contextualizing predictive minds. Neuroscience and Biobehavioral Reviews. 2025 1;168. doi:10.1016/j.neubiorev.2024.105948.

13. Gershman SJ, Blei DM, Niv Y. Context, Learning, and Extinction. Psychological Review. 2010;117:197–209. doi:10.1037/a0017808.

14. Schwoebel S, Marković D, Smolka MN, Kiebel SJ. Balancing control: A Bayesian interpretation of habitual and goal-directed behavior. Journal of Mathematical Psychology. 2021;100:102472. doi:10.1016/j.jmp.2020.102472.

15. Wilson RC, Takahashi YK, Schoenbaum G, Niv Y. Orbitofrontal Cortex as a Cognitive Map of Task Space. Neuron. 2014 1;81:267–79. doi:10.1016/J.NEURON.2013.11.005.

16. Hampton AN, Bossaerts P, O’Doherty JP. The role of the ventromedial prefrontal cortex in abstract state-based inference during decision making in humans. Journal of Neuroscience. 2006 8;26:8360–7. doi:10.1523/JNEUROSCI.1010-06.2006.

17. Maren S, Phan KL, Liberzon I. The contextual brain: Implications for fear conditioning, extinction and psychopathology. Nature Reviews Neuroscience. 2013 6;14:417–28. doi:10.1038/nrn3492.

18. Rao RPN, Ballard DH. Predictive coding in the visual cortex: a functional interpretation of some extra-classical receptive-field effects. Nature Neuroscience. 1999;2:79–87. doi:10.1038/4580.

19. Knill DC, Pouget A. The Bayesian brain: The role of uncertainty in neural coding and computation. Trends in Neurosciences. 2004;27:712–9. doi:10.1016/j.tins.2004.10.007.

20. Friston K. The free-energy principle: a unified brain theory? Nature reviews neuroscience. 2010;11(2):127–38. doi:10.1038/nrn2787.

21. Friston K, FitzGerald T, Rigoli F, Schwartenbeck P, Pezzulo G, et al. Active inference and learning. Neuroscience & Biobehavioral Reviews. 2016;68:862–79. doi:10.1016/j.neubiorev.2016.06.022.

22. Donoso M, Collins AGE, Koechlin E. Foundations of human reasoning in the prefrontal cortex. Science. 2014 6;344:1481–6. doi:10.1126/science.1252254.

23. Gershman SJ, Blei DM. A tutorial on Bayesian nonparametric models. Journal of Mathematical Psychology. 2012 2;56:1–12. doi:10.1016/j.jmp.2011.08.004.

24. Tse D, Langston RF, Kakeyama M, Bethus I, Spooner PA, Wood ER, et al. Schemas and Memory Consolidation. Science. 2007 4;316:76–82. doi:10.1126/science.1135935.

25. Tse D, Takeuchi T, Kakeyama M, Kajii Y, Okuno H, Tohyama C, et al. Schema-Dependent Gene Activation and Memory Encoding in Neocortex. Science. 2011 8;333:891–5. doi:10.1126/science.1207079.

26. Ghosh VE, Gilboa A. What is a memory schema? A historical perspective on current neuroscience literature. Neuropsychologia. 2014 1;53:104–14. doi:10.1016/j.neuropsychologia.2013.11.010.

27. Lieder F, Griffiths TL. Resource-rational analysis: Understanding human cognition as the optimal use of limited computational resources. Behavioral and Brain Sciences. 2019;43. doi:10.1017/S0140525X1900061X.

28. Bhui R, Lai L, Gershman SJ. Resource-rational decision making. Current Opinion in Behavioral Sciences. 2021 10;41:15–21. doi:10.1016/j.cobeha.2021.02.015.

29. Keramati M, Smittenaar P, Dolan RJ, Dayan P. Adaptive integration of habits into depth-limited planning defines a habitual-goal-directed spectrum. Proceedings of the National Academy of Sciences of the United States of America. 2016 11;113:12868–73. doi:10.1073/pnas.1609094113.

30. Wood W, Rünger D. Psychology of habit. Annual Review of Psychology. 2016;67:289–314. doi:10.1146/annurev-psych-122414-033417.

31. Gershman SJ. Habituation as optimal filtering. iScience. 2024 8;27. doi:10.1016/j.isci.2024.110523.

32. Schwöbel S, Marković D, Smolka MN, Kiebel S. Joint modeling of choices and reaction times based on Bayesian contextual behavioral control. PLoS Computational Biology. 2024 7;20. doi:10.1371/journal.pcbi.1012228.

33. Bein O, Niv Y. Schemas, reinforcement learning and the medial prefrontal cortex. Nature Reviews Neuroscience. 2025 3;26:141–57. doi:10.1038/s41583-024-00893-z.

34. Frigyik BA, Kapila A, Gupta MR, Uw UW. Introduction to the Dirichlet Distribution and Related Processes. UWEE Tech Report Series; 2010.

35. Neal RM. Bayesian Mixture Modeling. In: Bayesian Mixture Modeling. Dordrecht: Springer Netherlands; 1992. p. 197–211.

36. Ishwaran H, Zarepour M. Exact and Approximate Sum Representations for the Dirichlet Process. The Canadian Journal of Statistics. 2002 6;30:269–83. doi:https://www.jstor.org/stable/3315951.

37. Teh YW, Jordan MI, Beal MJ, Blei DM. Hierarchical Dirichlet Processes. Journal of the American Statistical Association. 2005;101:1–198.

38. Fox EB, Sudderth EB, Jordan MI, Willsky AS. A sticky HDP-HMM with application to speaker diarization. Annals of Applied Statistics. 2011 6;5:1020–56. doi:10.1214/10-AOAS395.

39. Attias H. Planning by probabilistic inference. In: International workshop on artificial intelligence and statistics. PMLR; 2003. p. 9–16.

40. Botvinick M, Toussaint M. Planning as inference. Trends in cognitive sciences. 2012;16(10):485–8. doi:10.1016/j.tics.2012.08.006.

41. Solway A, Botvinick MM. Goal-directed decision making as probabilistic inference: a computational framework and potential neural correlates. Psychological review. 2012;119(1):120–54. doi:10.1037/a0026435.

42. Bishop C. Pattern recognition and machine learning. vol. 4. New York: springer; 2006.

43. Friston K. The free-energy principle: a rough guide to the brain? Trends in cognitive sciences. 2009;13(7):293–301. doi:10.1016/j.tics.2009.04.005.

44. Da Costa L, Parr T, Sajid N, Veselic S, Neacsu V, Friston K. Active inference on discrete state-spaces: A synthesis. Journal of Mathematical Psychology. 2020;99:102447. doi:10.1016/j.jmp.2020.102447.

45. Sajid N, Ball PJ, Parr T, Friston KJ. Active Inference: Demystified and Compared. Neural Computation. 2021;712:1–39. doi:10.1162/neco_a_01357.

46. Friston K, Rigoli F, Ognibene D, Mathys C, Fitzgerald T, Pezzulo G. Active inference and epistemic value. Cognitive Neuroscience. 2015;6:187–214. doi:10.1080/17588928.2015.1020053.

47. Littman ML. A tutorial on partially observable Markov decision processes. Journal of Mathematical Psychology. 2009;53(3):119–25. doi:10.1016/j.jmp.2009.01.005.

48. Sutton RS, Barto AG. Reinforcement Learning: An Introduction. vol. 1. Cambridge: MIT press; 1998.

49. Yedidia JS, Freeman WT, Weiss Y. Understanding Belief Propagation and its Generalizations. Exploring artificial intelligence in the new millennium. 2003;8:236–9.

50. Schwöbel S, Kiebel S, Marković D. Active inference, belief propagation, and the bethe approximation. Neural computation. 2018;30(9):2530–67. doi:10.1162/neco_a_01108.

51. Erp BV, Nuijten WWL, Vries BD. Online Structure Learning with Dirichlet Processes through Message Passing. In: International Workshop on Active Inference. Springer Nature Switzerland; 2024. p. 91–104.

52. Thrailkill EA, Bouton ME. Contextual control of instrumental actions and habits. Journal of Experimental Psychology: Animal Learning and Cognition. 2015 1;41:69–80. doi:10.1037/xan0000045.

53. Corbit LH, Balleine BW. The role of prelimbic cortex in instrumental conditioning. Behavioural Brain Research. 2003;146:145–57. doi:10.1016/j.bbr.2003.09.023.

54. Cools R, Clark L, Owen AM, Robbins TW. Defining the Neural Mechanisms of Probabilistic Reversal Learning Using Event-Related Functional Magnetic Resonance Imaging. The Journal of Neuroscience. 2002 6;22:4563–7. doi:10.1523/JNEUROSCI.22-11-04563.2002.

55. Frank MJ, Seeberger LC, O’Reilly RC. By carrot or by stick: Cognitive reinforcement learning in Parkinsonism. Science. 2004 12;306:1940–3. doi:10.1126/science.1102941.

56. Behrens TEJ, Woolrich MW, Walton ME, Rushworth MFS. Learning the value of information in an uncertain world. Nature Neuroscience. 2007 9;10:1214–21. doi:10.1038/nn1954.

57. Wagner BJ, Wolf HB, Kiebel SJ. Action repetition biases choice in context-dependent decision-making. Communications Psychology. 2025. doi:10.1038/s44271-025-00363-x.

58. Marković D, Stojić H, Schwöbel S, Kiebel SJ. An empirical evaluation of active inference in multi-armed bandits. Neural Networks. 2021 12;144:229–46. doi:10.1016/j.neunet.2021.08.018.

59. Collins A, Koechlin E. Reasoning, learning, and creativity: Frontal lobe function and human decision-making. PLoS Biology. 2012 3;10. doi:10.1371/journal.pbio.1001293.

60. Todd TP, Winterbauer NE, Bouton ME. Contextual control of appetite. Renewal of inhibited food-seeking behavior in sated rats after extinction. Appetite. 2012 4;58:484–9. doi:10.1016/j.appet.2011.12.006.

61. Bouton ME, Todd TP. A fundamental role for context in instrumental learning and extinction. Behavioural Processes. 2014;104:91–8. doi:10.1016/j.beproc.2014.02.012.

62. Adams CD. Variations in the sensitivity of instrumental responding to reinforcer devaluation. The Quarterly Journal of Experimental Psychology Section B. 1982 5;34:77–98. doi:10.1080/14640748208400878.

63. Dickinson A. Omission Learning after Instrumental Pretraining. The Quarterly Journal of Experimental Psychology: Section B. 1998;51:271–86. Available from: 10.1080/713932679. doi:10.1080/713932679.

64. Feldman J. The Simplicity Principle in Human Concept Learning. Current directions in psychological science. 2003;12:227–32. doi:10.1046/j.0963-7214.2003.01267.x.

65. Briscoe E, Feldman J. Conceptual complexity and the bias/variance tradeoff. Cognition. 2011 1;118:2–16. doi:10.1016/j.cognition.2010.10.004.

66. Yin HH, Knowlton BJ. The role of the basal ganglia in habit formation. Nature Reviews Neuroscience. 2006;7:464–76. doi:10.1038/nrn1919.

67. Ricker ST, Bouton ME. Reacquisition following extinction in appetitive conditioning. Animal Learning and Behavior. 1996;24:423–36. doi:10.3758/BF03199014.

68. Napier RM, Macrae M, Kehoe EJ. Rapid Reacquisition in Conditioning of the Rabbit’s Nictitating Membrane Response. Journal of Experimental Psychology: Animal Behavior Processes. 1992;18:182–92. doi:10.1037/0097-7403.18.2.182.

69. Bouton ME, Bolles RC. Role of conditioned contextual stimuli in reinstatement of extinguished fear. Journal of Experimental Psychology: Animal Behavior Processes. 1979;5:368–78. doi:10.1037/0097-7403.5.4.368.

70. Rescorla RA, Heth CD. Reinstatement of fear to an extinguished conditioned stimulus. Journal of Experimental Psychology: Animal Behavior Processes. 1975 1;1:88–96. doi:10.1037/0097-7403.1.1.88.

71. Gilboa A, Marlatte H. Neurobiology of Schemas and Schema-Mediated Memory. Trends in Cognitive Sciences. 2017 8;21:618–31. doi:10.1016/j.tics.2017.04.013.

72. Santoro A, Frankland PW, Richards BA. Memory transformation enhances reinforcement learning in dynamic environments. Journal of Neuroscience. 2016 11;36:12228–42. doi:10.1523/JNEUROSCI.0763-16.2016.

73. Schuck NW, Cai MB, Wilson RC, Niv Y. Human Orbitofrontal Cortex Represents a Cognitive Map of State Space. Neuron. 2016 9;91:1402–12. doi:10.1016/j.neuron.2016.08.019.

74. Sharpe MJ, Stalnaker T, Schuck NW, Killcross S, Schoenbaum G, Niv Y. An Integrated Model of Action Selection: Distinct Modes of Cortical Control of Striatal Decision Making. Annual Review of Psychology. 2019 1;70:53–76. doi:10.1146/annurev-psych-010418-102824.

75. Chan SCY, Niv Y, Norman KA. A probability distribution over latent causes, in the orbitofrontal cortex. Journal of Neuroscience. 2016 7;36:7817–28. doi:10.1523/JNEUROSCI.0659-16.2016.

76. Saez A, Rigotti M, Ostojic S, Fusi S, Salzman CD. Abstract Context Representations in Primate Amygdala and Prefrontal Cortex. Neuron. 2015 8;87:869–81. doi:10.1016/j.neuron.2015.07.024.

77. Rosenbaum RS, Gilboa A, Moscovitch M. Case studies continue to illuminate the cognitive neuroscience of memory. Annals of the New York Academy of Sciences. 2014;1316:105–33. doi:10.1111/nyas.12467.

78. Gilboa A, Moscovitch M. Ventromedial prefrontal cortex generates pre-stimulus theta coherence desynchronization: A schema instantiation hypothesis. Cortex. 2017 2;87:16–30. doi:10.1016/J.CORTEX.2016.10.008.

79. Frühholz S, Godde B, Lewicki P, Herzmann C, Herrmann M. Face recognition under ambiguous visual stimulation: fMRI correlates of “encoding styles”. Human Brain Mapping. 2011 10;32:1750–61. doi:10.1002/HBM.21144.

80. Wikenheiser AM, Schoenbaum G. Over the river, through the woods: cognitive maps in the hippocampus and orbitofrontal cortex. Nature Reviews Neuroscience 2016 17:8. 2016 6;17:513–23. doi:10.1038/nrn.2016.56.

81. Behrens TEJ, Muller TH, Whittington JCR, Mark S, Baram AB, Stachenfeld KL, et al. What Is a Cognitive Map? Organizing Knowledge for Flexible Behavior. Neuron. 2018 10;100:490–509. doi:10.1016/J.NEURON.2018.10.002.

82. Knudsen EB, Wallis JD. Taking stock of value in the orbitofrontal cortex. Nature Reviews Neuroscience 2022 23:7. 2022 4;23:428–38. doi:10.1038/s41583-022-00589-2.

83. Padoa-Schioppa C, Conen KE. Orbitofrontal Cortex: A Neural Circuit for Economic Decisions. Neuron. 2017 11;96:736–54. doi:10.1016/J.NEURON.2017.09.031.

84. Zhou J, Gardner MP, Schoenbaum G. Is the core function of orbitofrontal cortex to signal values or make predictions? Current opinion in behavioral sciences. 2021 10;41:1–9. doi:10.1016/J.COBEHA.2021.02.011.

85. Klein-Flügge MC, Bongioanni A, Rushworth MFS. Medial and orbital frontal cortex in decision-making and flexible behavior. Neuron. 2022 9;110:2743–70. doi:10.1016/J.NEURON.2022.05.022.

86. Moneta N, Grossman S, Schuck NW. Representational spaces in orbitofrontal and ventromedial prefrontal cortex: task states, values, and beyond. Trends in Neurosciences. 2024 12;47:1055–69. doi:10.1016/J.TINS.2024.10.005.

87. Everitt BJ, Robbins TW. Drug addiction: Updating actions to habits to compulsions ten years on. Annual Review of Psychology. 2016;67:23–50. doi:10.1146/annurev-psych-122414-033457.

88. Marchant NJ, Kaganovsky K, Shaham Y, Bossert JM. Role of corticostriatal circuits in context-induced reinstatement of drug seeking. Brain Research. 2015 12;1628:219–32. doi:10.1016/j.brainres.2014.09.004.

89. Heinz A, Kiefer F, Smolka MN, Endrass T, Beste C, Beck A, et al. Addiction Research Consortium: Losing and regaining control over drug intake (ReCoDe)—From trajectories to mechanisms and interventions. Addiction Biology. 2020 3;25. doi:10.1111/adb.12866.

90. Spanagel R, Bach P, Banaschewski T, Beck A, Bermpohl F, Bernardi RE, et al. The ReCoDe addiction research consortium: Losing and regaining control over drug intake—Findings and future perspectives. Addiction Biology. 2024 7;29. doi:10.1111/adb.13419.

91. Smith R, Schwartenbeck P, Parr T, Friston KJ. An Active Inference Approach to Modeling Structure Learning: Concept Learning as an Example Case. Frontiers in Computational Neuroscience. 2020 5;14. doi:10.3389/fncom.2020.00041.

92. Inostroza M, Born J. Sleep for preserving and transforming episodic memory. Annual Review of Neuroscience. 2013 7;36:79–102. doi:10.1146/annurev-neuro-062012-170429.

93. Lewis PA, Durrant SJ. Overlapping memory replay during sleep builds cognitive schemata. Trends in Cognitive Sciences. 2011 8;15:343–51. doi:10.1016/J.TICS.2011.06.004.

